# A high-resolution view of RNA endonuclease cleavage in *Bacillus subtilis*

**DOI:** 10.1101/2023.03.12.532304

**Authors:** James C. Taggart, Julia Dierksheide, Hannah LeBlanc, Jean-Benoît Lalanne, Sylvain Durand, Frédérique Braun, Ciarán Condon, Gene-Wei Li

**Affiliations:** Department of Biology, Massachusetts Institute of Technology, Cambridge, MA 02142, USA; Department of Physics, Massachusetts Institute of Technology, Cambridge, MA 02142, USA; Expression Génétique Microbienne (EGM), CNRS, Université Paris Cité, Institut de Biologie Physico-Chimique, 13 rue Pierre et Marie Curie, 75005 Paris, France; Current address: Department of Systems Biology, Harvard Medical School, Boston, MA, USA; Current address: Department of Genome Sciences, University of Washington, Seattle, WA, USA.

## Abstract

RNA endonucleases are the rate-limiting initiator of decay for many bacterial mRNAs. However, the positions of cleavage and their sequence determinants remain elusive even for the well-studied *Bacillus subtilis*. Here we present two complementary approaches – transcriptome-wide mapping of endoribonucleolytic activity and deep mutational scanning of RNA cleavage sites – that reveal distinct rules governing the specificity among *B. subtilis* endoribonucleases. Detection of RNA terminal nucleotides in both 5′- and 3′-exonuclease-deficient cells revealed >10^3^ putative endonucleolytic cleavage sites with single-nucleotide resolution. We found a surprisingly weak consensus for RNase Y targets, a contrastingly strong primary sequence motif for EndoA targets, and long-range intramolecular secondary structures for RNase III targets. Deep mutational analysis of RNase Y cleavage sites showed that the specificity is governed by many disjointed sequence features, each with mild contributions. Our results highlight the delocalized nature of mRNA stability determinants and provide a strategy for elucidating endoribonuclease specificity *in vivo*.

## INTRODUCTION

Predicting protein abundance from an arbitrary gene sequence remains an aspirational goal for quantitative biology. Decades of research has explored the encoding of rates of transcription and translation initiation (Taggart et al., 2021), resulting in predictive models for how RNA polymerase and the ribosome recognize promoters and ribosome binding sites (Brewster et al., 2012; Salis, 2011; Salis et al., 2009; Urtecho et al., 2019). Comparatively poorly characterized, however, is post-transcriptional control of RNA abundance by ribonucleases, factors which exert widespread quantitative control on bacterial gene expression levels (Chen et al., 2015; Khemici et al., 2015; Moffitt et al., 2016). Understanding the encoding of ribonuclease cleavage sites in bacterial mRNA sequences is therefore a critical question in quantitative biology.

In the Gram-positive bacterium *Bacillus subtilis*, cleavage by an endoribonuclease is thought to initiate the decay of a large fraction of cellular mRNAs (Durand et al., 2012a; Lehnik-Habrink et al., 2011a, 2012; Shahbabian et al., 2009). The cleaved RNA fragments typically have a 5′-phosphate and a 3′-hydroxyl group, which are subsequently degraded by a 5′-3′ exoribonuclease (RNase J1) and by a combination of 3′-5′ exoribonucleases (PNPase, RNase R, RNase PH, and YhaM) (Trinquier et al., 2020), respectively. The endoribonuclease RNase Y appears to drive the turnover of the majority of mRNAs in *B. subtilis*, and its depletion results in the accumulation of hundreds of transcripts (Durand et al., 2012a; Lehnik-Habrink et al., 2011a; Shahbabian et al., 2009). For several well-studied mRNAs, RNase Y cleavage is specific to certain positions and can drive operon mRNA maturation (Braun et al., 2017; Commichau et al., 2009; DeLoughery et al., 2018; Noone et al., 2014). Unfortunately, though RNase Y has been identified as a key player shaping the transcriptomes of Gram-positive bacteria, our understanding of which mRNA sequences *B. subtilis* RNase Y cleaves remains relatively limited. Studies have suggested *B. subtilis* RNase Y recognizes single stranded RNA, is stimulated by downstream secondary structure, and has a preference for adenosine and uridine-rich sequence (Braun et al., 2017; Shahbabian et al., 2009). Orthologs of RNase Y in *Staphylococcus aureus* and *Streptococcus pyogenes*, organisms for which RNase Y is not essential and appears to play a more limited role compared to *B. subtilis*, additionally demonstrate a preference for cleavage downstream of a guanosine (Khemici et al., 2015) (Benda et al., 2021). However, these limited sequence preferences cannot explain the highly specific locations of cleavage that have been observed for the *B. subtilis* RNase Y (DeLoughery et al., 2018).

In addition to RNase Y, several other endoribonucleases contribute to mRNA decay. RNase III has been suggested to cleave and initiate decay for dozens of mRNAs in *B. subtilis* (DiChiara et al., 2016). Although the *E. coli* RNase III strictly cleaves double-stranded RNAs, the *B. subtilis* RNase III has been implicated to deviate from this rule in a way that remains unknown (DiChiara et al., 2016). Other minor endoribonucleases, such as the RNase toxin EndoA (a homolog of *E. coli* MazF), may also initiate mRNA decay and produce different 5′ and 3′ moieties (Cerullo et al., 2022; Ingle et al., 2022; Leroy et al., 2017). Their positions of cleavage and subsequent decay processes remain poorly described (Pellegrini et al., 2005).

A number of studies have mapped the target repertoire of major bacterial endoribonucleases (Altuvia et al., 2018; Broglia et al., 2020; Chao et al., 2017; DiChiara et al., 2016; Gordon et al., 2017; Khemici et al., 2015). These measurements detect the terminal nucleotides of RNA decay intermediates and analyze their abundance upon endoribonuclease inactivation. The results are confounded by several factors. First, because cleaved decay intermediates are typically unstable, the detected end signals are often close to background levels and may be biased by their own stability. Second, the sequential actions of several endoribonucleases and/or exoribonucleases on the same mRNA may result in RNA ends that are shifted from the initial position that is cleaved by the endoribonuclease under investigation. As a result, low-throughput approaches are often required to precisely determine exact positions of cleavage for each target (Braun et al., 2017; Bruscella et al., 2011; DiChiara et al., 2016; Yao and Bechhofer, 2010).

In this work, we first present a global approach to map bacterial endoribonuclease cleavage sites by combining high-resolution RNA end-mapping with genetic perturbation of all known exonucleases participating in *B. subtilis* mRNA decay. By stabilizing mRNA decay intermediates in exonuclease deficient *B. subtilis* cells, this “stabilized end-sequencing” approach revealed thousands of putative positions of endoribonuclease cleavage and a previously unknown nuclease activity targeting RNA 5′ ends. Furthermore, using a panel of endoribonuclease knockouts, we dramatically expanded the known target repertoire for RNase Y, RNase III, and EndoA. Our results captured a revised EndoA cleavage motif, elucidated long-range RNA folds that dictate RNase III cleavage, and a surprisingly weak consensus among RNase Y cleavage sites, despite their high specificity. To further investigate the sequence determinants for RNase Y cleavage, we developed a deep mutational scanning approach to interrogate individual cleavage sites. Interestingly, the residues that affect cleavage activity are scattered across a region over 40 nucleotides long. These results suggest that the highly specific RNase Y cleavage does not follow the conventional model of protein-nucleic acid interactions mediated by compact sequence or structural motifs. Instead, the “cleavage code” is specified by a collection of weak interactions embedded in an extended region.

## RESULTS

### Stabilized end-sequencing reveals positions of endoribonucleolytic activity

In the absence of additional nucleolytic activity, one can map the positions of cleavage by an endoribonuclease by simply observing the termini of its cleavage products. *In vivo*, however, exonucleases typically rapidly act on endonuclease cleavage products, rendering the original RNA endonuclease cleavage sites difficult to detect. If all exoribonucleolytic activity targeting either 3′ or 5′ ends is ablated, endoribonuclease cleavage at a particular site should result in stably accumulating products whose ends correspond to the position of cleavage (Figure 1A). Strains combining knockouts of all known exonucleases in either the 5′ or 3′ direction are viable in *B. subtilis*, meaning such a measurement is theoretically possible in this species (Figaro et al., 2013; Oussenko et al., 2005). If RNA ends are separately mapped in both a knockout of RNase J1, the only known 5′-3′ exonuclease, and a knockout of all four known 3′-5′ exonucleases (a knockout of *yhaM, rph, pnpA,* and *rnr*, hereafter referred to as a 4-exo knockout), exact positions of endoribonucleolytic activity can be captured as pairs of adjacent 3′ and 5′ ends that appear across the two strains.

**Figure 1.**
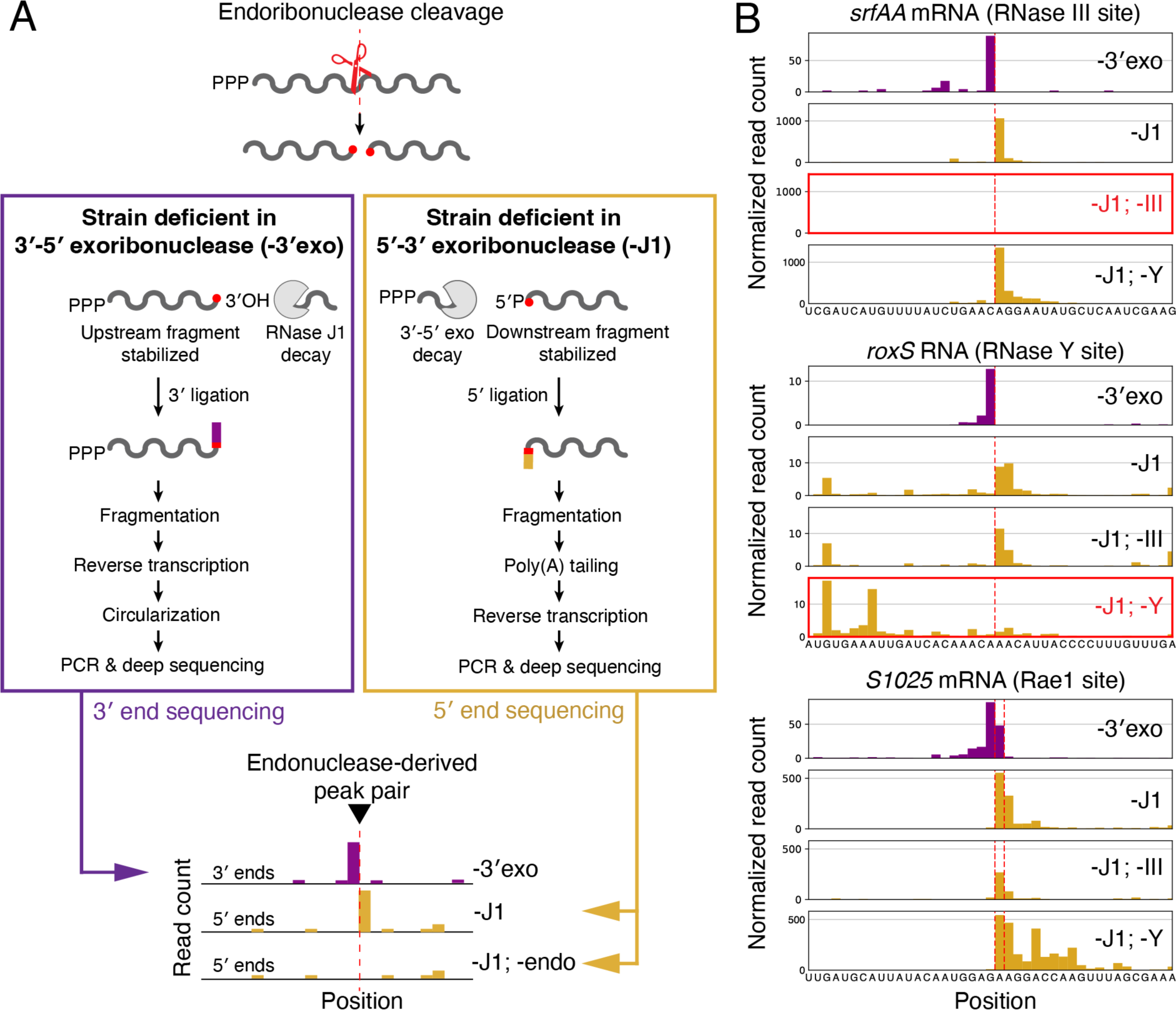
Workflow and validation of endoribonuclease cleavage mapping approach. (A) Deletion of exoribonucleases results in stable accumulation of RNA decay intermediates. A schematic mRNA is shown with endoribonuclease cleavage (red scissors) occurring, with remaining exoribonucleases indicated in gray. RNAs are shown with 5′-P and 3′-OH characteristic of the most common endoribonucleases in *B. subtilis*. Newly generated RNA ends can be captured through RNA ligation allowing mapping of cleavage positions by 3′ and 5′ end sequencing. Adapters used to capture RNA ends indicated in purple for 3′ end sequencing and yellow for 5′ end sequencing, with matching color scheme in schematic data below. (B) Validation of endonuclease cleavage site detection using known positions of endoribonucleolytic cleavage by RNase III, RNase Y, and Rae1 (DiChiara et al., 2016; Durand et al., 2015; Leroy et al., 2017). Yellow indicates 5′ end sequencing data and purple indicates 3′ end sequencing data. Red dotted line represents manually annotated cleavage positions. Plotted are reads per million CDS-mapping reads, normalized to the average 3′-mapped Rend-seq RPM in this window.

Based on this principle, we implemented a “stabilized end-sequencing” approach to create a transcriptome-wide map of endoribonuclease activity by capturing the 3′ hydroxyl and 5′ monophosphate groups that are generated by endonucleases such as RNases Y and III (the primary endonucleases targeting mRNAs in *B. subtilis*) (Figure 1A; (Bechhofer and Deutscher, 2019; Leroy et al., 2017)). To quantitatively capture 3′ hydroxyl groups, we utilized a previously established method for RNA 3′ ligation and deep sequencing, hereafter referred to as 3′ end sequencing (Figure 1A, left; (Dar et al., 2016; Herzel et al., 2022; Mandell et al., 2021)). To complement 3′ end mapping, we specifically captured 5′ monophosphates through T4 RNA Ligase 1 treatment, allowing quantification of cleavage-derived 5′ ends by deep sequencing, hereafter referred to as 5′ end sequencing (Figure 1A, right). By performing 3′ end sequencing on a 4-exo knockout (strain CCB396) and 5′ end sequencing on an RNase J1 knockout (strain CCB434), we captured 3′/5′ end pairs derived from endoribonuclease activity transcriptome-wide. Combining these exoribonuclease perturbations with endoribonuclease knockouts, we can attribute these cleavage signatures to specific enzymes. End-enriched RNA sequencing (Rend-seq) can additionally be performed to quantify expression of putative endoribonuclease targets and provide an orthogonal RNA end-mapping approach (Lalanne et al., 2018).

This stabilized end-sequencing strategy robustly captures positions of previously documented endoribonucleolytic cleavage sites. Among 13 previously characterized sites cleaved by either RNase Y, RNase III, or Rae1 that were individually detected (Braun et al., 2017; Bruscella et al., 2011; DiChiara et al., 2016; Durand et al., 2015; Leroy et al., 2017; Noone et al., 2014; Yao and Bechhofer, 2010), we identified at least one clear 3′/5′ pair at or near the annotated cleavage position (Figure 1B, Figure S1). 5′ end sequencing of RNase Y+J1 or III+J1 double mutant strains (CCB760 and BG879) confirmed that we can detect cleavage by both endoribonucleases and accurately assign the enzymes responsible for cleavage (Figure 1B).

Importantly, the paired end-mapping strategy provides clear advantages over measurements of a single end type. Because we capture both RNA ends generated by endoribonuclease activity, multiple adjacent positions of cleavage can be resolved, as appears to be the case for the first characterized substrate of Rae1, *S1025* (Figure 1B, bottom). Without these two paired channels of information, it is difficult to determine whether signal at adjacent positions is derived from multiple positions of endoribonucleolytic cleavage or residual exoribonuclease activity by unknown enzymes, the latter of which may be occurring after endoribonucleolytic cleavage of the RoxS RNA (Figure 1B, middle). Precisely resolving cleavage positions becomes particularly relevant in cases such as the endoribonucleolytic cleavages observed in *hbs*, where many nearby positions are known to be subject to RNase Y cleavage (Figure S1, (Braun et al., 2017)).

Using these data, we first identified putative positions of cleavage within transcripts whose abundance is known to be regulated by RNA endonuclease activity. Recent work has shown that the RNA decay machinery in *B. subtilis* is subject to extensive autoregulation and cross-regulation, suggesting that cleavage sites should be detectable within the RNAs encoding decay-associated proteins (DeLoughery et al., 2018; DiChiara et al., 2016; Korobeinikova et al., 2023). This expectation is borne out in our stabilized end-sequencing, where multiple putative positions of endoribonucleolytic cleavage are detected within the transcripts encoding RNases Y, III, and the Y-complex, a complex of three small proteins (YlbF, YmcA, and YaaT) that modulates RNase Y activity (Deloughery et al., 2016) (Figure 2, Figure S2). Further, a number of additional transcripts, including that encoding sporulation master regulator SinR, have been experimentally characterized as stabilized upon deletion of RNase Y (Deloughery et al., 2016; Lehnik-Habrink et al., 2011a). Paired peaks are clearly visible within each of these transcripts (Fig 2, Fig S2). Taken together, our ability to robustly identify known and previously unknown positions of cleavage suggests that stabilized end-sequencing provide a high-resolution, genome-wide view of RNA endonuclease activity in *B. subtilis*.

**Figure 2.**
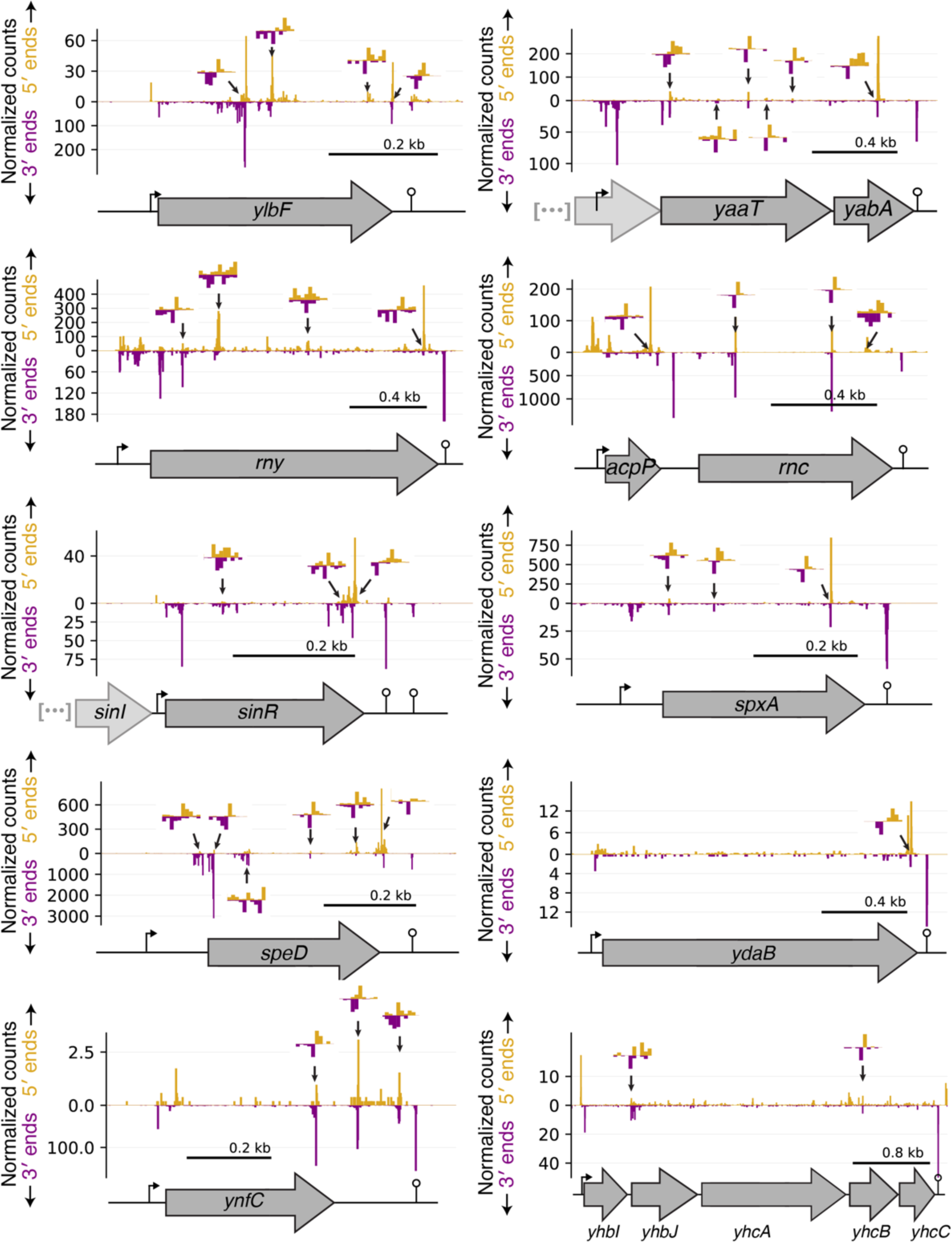
Cleavage within transcripts known to be destabilized by endoribonucleolytic cleavage. 5′ and 3′ end sequencing data are shown in yellow and purple, respectively. Plotted are reads per million CDS-mapping reads. Manually annotated putative cleavage sites (positions of adjacent 3′ and 5′ read density) are highlighted with arrows. Insets show highlighted sites at single-nucleotide resolution, with the Y-axis in each direction rescaled to the maximal value within the inset region. Data are aligned to a schematic representation of the relevant genomic locus. Annotated promoter and transcriptional terminators, indicated by bent arrows and lollipops, respectively, were identified using Rend-seq in SSB1002. Sequence context and RNase Y dependence of highlighted sites shown in Figure S2. Consistent with a recent report (Korobeinikova et al., 2023), multiple 5′ and 3′ end signals were observed in the 5′ UTR of *rny*.

### Systematic identification of positions of endoribonuclease cleavage

To systematically identify putative positions of endonuclease cleavage, we implemented a computational approach to call abutting pairs of peaks in 3′ and 5′ end sequencing across our 4-exo and RNase J1 knockout data, respectively (Figure S3A). To do so, we assessed the signal at each position relative to the average in a local window, calling positions exceeding a signal-to-background threshold as peaks. Positions for which a peak is called in both 5′ and 3′ end sequencing with a 1-nucleotide offset (a 5′ peak downstream of a 3′ peak) are called as peak pairs and represent putative sites of endoribonucleolytic activity. This approach reveals 1798 putative positions of cleavage in mRNAs. It should be noted that we are unable to identify cleavage events proximal to the ends of transcription units, because short RNA decay fragments (<35 nt) are excluded during 5′ end sequencing library preparation (see Methods).

To assign putative cleavage positions to the activity of specific endoribonucleases, we identified the positions that have diminished end-signals when an endoribonuclease is knocked out (Fig S3B). To account for changes in expression levels in strains with and without the endoribonuclease, we first normalized end-sequencing signals to local RNA abundance measured by Rend-seq in the same strain. We then quantified the sensitivity of the expression-normalized signal to endoribonuclease knockout, by computing the ratio of 5′-end signals (in the absence of RNase J1) between strains with and without the endoribonuclease (Materials and Methods). This “sensitivity” score will be elevated for positions cleaved by the corresponding endoribonuclease, as was previously shown by DiChiara et al. (DiChiara et al., 2016).

### *B. subtilis* RNase III cleaves within long-range intramolecular structures

We first characterized the cleavage profile of RNase III, motivated by previous reports of unexpected cleavage patterns in *B. subtilis* (DiChiara et al., 2016). RNase III plays well-characterized roles in the processing of noncoding RNAs (DiChiara et al., 2016; Herskovitz and Bechhofer, 2000; Oguro et al., 1998) and the regulation of mRNAs (Durand et al., 2012b; Jahn and Brantl, 2013; Meißner et al., 2016). Several potential mRNA targets of this enzyme have previously been proposed, but only a small number of cleavage sites in these mRNAs have been identified (DiChiara et al., 2016; Gordon et al., 2017). Intriguingly, such cleavage sites often did not appear as pairs at a defined 2-nt spacing within RNA duplexes, contrasting with the hallmark of RNase III activity in *E. coli* (Altuvia et al., 2018). We reasoned that a more systematic, high-resolution mapping of RNase III cleavage sites in *B. subtilis* will help resolve this issue.

Our stabilized end-sequencing data revealed 68 RNase III cleavage sites located on 42 mRNAs (Table S3). These sites constitute a small fraction of the putative endonuclease cleavage sites we identified (5%) in the background strain (phage-cured (Gilet et al., 2015)), consistent with the notion that RNase III targets a small subset of mRNAs (Fig. 3B). Despite the sparseness of these sites, they are often present in the same mRNAs as other RNase III sites (45 of 68) (Figure 3A, Figure S1, and Table S3). These results suggest that the *B. subtilis* RNase III acts in a manner similar to the *E. coli* RNase III in cutting a pair of positions in RNA duplexes.

**Figure 3.**
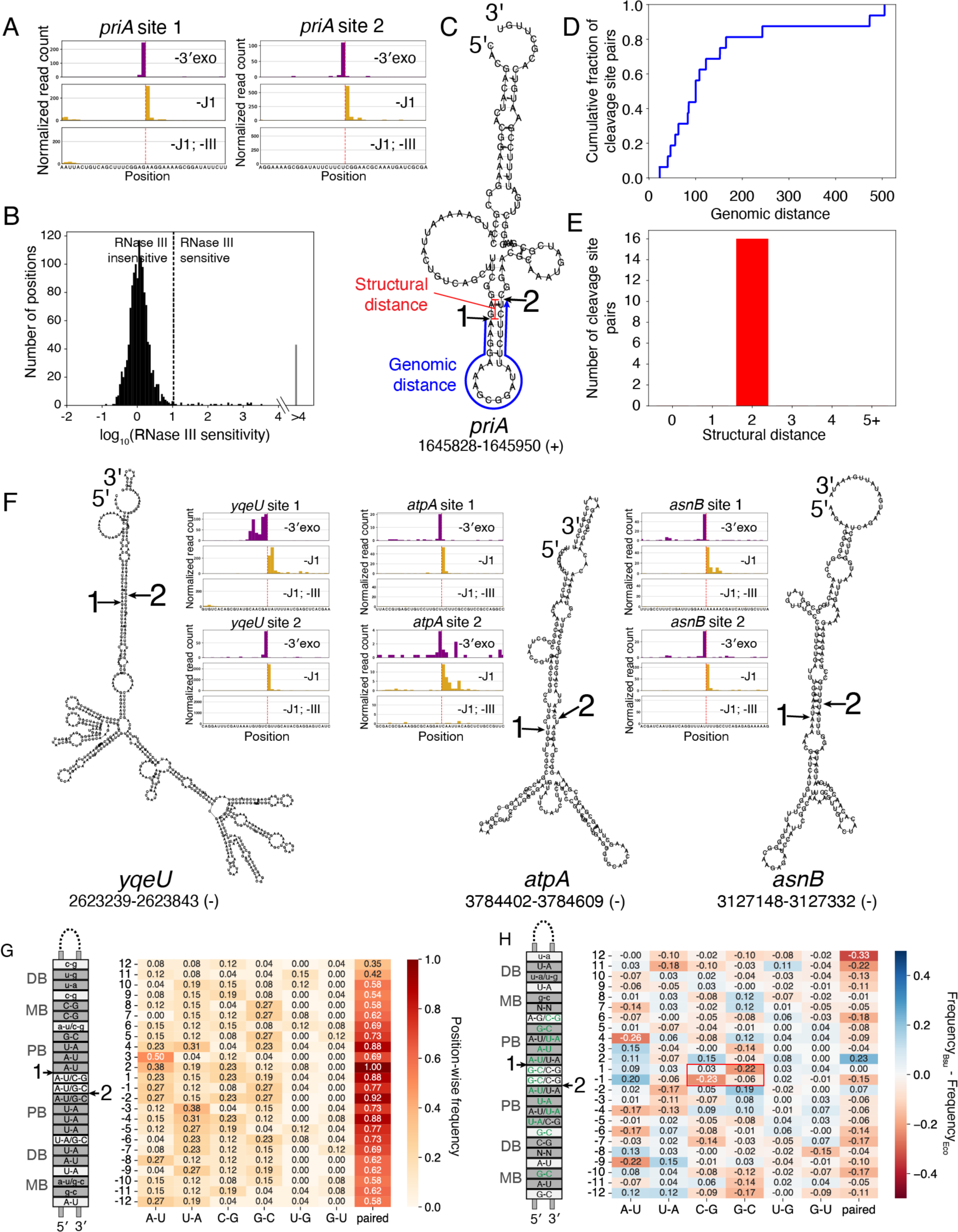
*B. subtilis* RNase III cleaves within long-range intramolecular secondary structures. (A) Identification of RNase III cleavage positions in the *priA* mRNA. 5′ and 3′ end sequencing data are shown in yellow and purple, respectively. Identified positions of cleavage are highlighted in red. Plotted are reads per million CDS-mapping reads, normalized to the average 3′-mapped Rend-seq RPM in this window. (B) Results of systematic identification of RNase III sites. Histogram shows distribution of endonuclease sensitivities for called peak pairs in 3′/5′ end sequencing of exoribonuclease knockouts (see Figure S3 and Methods), with black dashed line indicating threshold for calling dependence on RNase III. 68 sites exceeded this defined threshold. For 43 sites we were unable to calculate a sensitivity score due to an absence of 5′ end sequencing counts in our knockout. These sites are called as RNase III sensitive and are counted within the “>4” bin of the histogram. (C) Predicted secondary structure of the sequence surrounding the identified RNase III sites within the *priA* mRNA. Positions of cleavage are indicated with red arrows, with numbers corresponding to those in (A). Structural distance between these cleavage positions, defined as the defined as the length of the 3′ overhang generated by the cleavages, is annotated in red. Genomic distance, defined as the distance between the cleavage sites in the primary sequence, is annotated in blue. (D) Genomic distance (as defined in C) between pairs of unambiguous RNase III cleavage positions identified within 1 kb of one another and predicted to fall on opposite sides of an RNA stem with a 2 bp spacing between cleavage positions (32 of 68 positions). (E) Structural distance (as defined in C) generated by RNase III cleavage at both positions within the pairs of cleavage positions plotted in (D). (F) Predicted secondary structures of the sequence surrounding the identified RNase III sites within the mRNAs encoding the *yqeU*, *atpA*, and *asnB* genes. Insets are plotted as in (A). (G) Consensus sequence-structure motif of RNase III targets in *B. subtilis*. The predicted structures for each pair of RNase III-sensitive cleavage sites that generate a 2-nt 3′ overhang were aligned to the positions of cleavage, “1” and “2”. The position-wise frequency of each type of base-pairing interaction is shown. Note that the base pairing frequencies sum to the fraction of sites paired at each position. The most frequent base pair at each position is shown in the schematic on the left, with lower-case characters used to designate positions for which the most frequent base pair occurs <20% of the time. When two base pair frequencies are within 0.02 of one another, both are separated by a slash (e.g., A-U/G-C). The gray regions labeled “PB”, “MB”, and “DB” correspond to the proximal, middle, and distal boxes (Gan et al., 2006; Zhang and Nicholson, 1997). (H) Comparison of base pairing frequences between *B. subtilis* and *E. coli* (Altuvia et al., 2018). The consensus *E. coli* sequence-structure motif is shown on the left with top base pairs at each position which are shared between *B. subtilis* and *E. coli* highlighted in green. Red box highlights the decrease in C-G or G-C pairing in *B. subtilis* relative to *E. coli*.

Indeed, of the 44 of these co-occurring cleavage sites that occur within 1 kb of each other, a large majority (36 of 44) are predicted to reside in RNA secondary structures. The extent of secondary structures is substantial, with up-to 505 nt between cleavage sites (Figure 3D). Despite the large contour length, the cleavage positions are almost always (32 of 36) separated by 2 base pairs on the opposite sides of RNA duplexes, which range in length from 10 to 36 basepairs (Figure 3C E, and F). 4 of the 36 co-occurring sites are cleaved at 2 adjacent positions, which may still result from the 2-bp rule on individual RNA molecules but are difficult to resolve in our ensemble measurement. It is possible that the minority of sites that do not form a clear intra-molecular duplex (32 of 68) participate in inter-molecular pairing, but the presence of such interactions remains to be determined.

We next sought to determine what sequence features within these specific secondary structures might be recognized by RNase III. To increase the number of RNase III sites used in this analysis, we included putative cleavage positions with a lower threshold on 5′ end sequencing depth (see Methods for details, Tables S5,6 for a list of putative cleavage positions with this relaxed threshold, and Figures S3C,E for endonuclease sensitivity distributions), restricting our analysis only to pairs of such putative cleavage positions that were predicted to pair with 2-bp spacing. Using this approach, we identified an additional 10 pairs of RNase III cleavage sites for a total of 26 pairs. The genomic distance between these sites is similar to those initially identified (Figure S3D).

From these pairs of RNase III cleavage sites, we calculated the position-wise sequence content and along the predicted RNA duplex, as well as the fraction of sites for which each position is paired, using the two cleavage positions as our reference points (Figure 3G). Through this, we generated a *B. subtilis* RNase III sequence-structure motif as has previously been reported for *E. coli* (Altuvia et al., 2018, Figure 3H). Like the *E. coli* homolog, the positions surrounding *B. subtilis* RNase III cleavage sites appear to be consistently paired, with the probability of pairing increasing towards the cleavage positions (reaching >90% at the -2/+2 positions) (Figure 3G). Similarly, a number of sequence features appear to be recognized similarly by these enzymes, including position-specific preferences for A-U and G-C base-pairing within the proximal box (PB) and an increased occurrence of U-G wobble pairing within the distal box (DB) (Altuvia et al., 2018; Pertzev and Nicholson, 2006). At each position from -6 to 6, at least one of the maximum-frequency base pairs is conserved between these species (Figure 3H, green text). In contrast to *E. coli*, however, the base pairs between cleavage positions do not demonstrate a clear preference for C-G/G-C pairing in our data (Altuvia et al., 2018) (Figure 3H, red box).

Together, these findings suggest that the *B. subtilis* RNase III displays similar substrate preferences to its evolutionarily distant *E. coli* homolog (Altuvia et al., 2018; Pertzev and Nicholson, 2006). Furthermore, the presence of extensive long-range RNA secondary structure for RNase III targets suggests that mRNA base pairing across hundreds of nucleotides is common in this organism and can directly influence gene expression regulation (Braun et al., 2017; Ruiz de los Mozos et al., 2013).

### Induction of EndoA activity in exoribonuclease knockouts reveals additional 5′ end trimming activity in *B. subtilis*

To our surprise, manual inspection of our Rend-seq data in all strains with an RNase J1 or 4-exo knockout showed prominent peaks across the transcriptome within the motif U/ACAUA (/ denotes the cleavage position, Figure 4A, B). This sequence resembles an extended version of the canonical U/ACAU cleavage motif of the endonuclease EndoA, an RNA interferase of the MazF family associated with the EndoA-EndoAI type II toxin-antitoxin system (Park et al., 2011; Pellegrini et al., 2005). The presence of adjacent 3′ and 5′ ends in Rend-seq data separating the first U/A dinucleotide within this motif strongly suggests endoribonuclease activity (Figure 4A, B), possibly through activation of this toxin in the exoribonuclease deficient strains used in this study. This U/ACAUA-specific cleavage is not observed in Rend-seq data from wild-type *B. subtilis* (Figure S4A). Consistent with EndoA-mediated cleavage, which leaves a 5′ hydroxyl group (Bechhofer and Deutscher, 2019; Pellegrini et al., 2005), we did not observe corresponding signals in our 5′ end sequencing data, which is specific to 5′ monophosphate groups, shown for a representative site in the *pgk* gene (Figure 4E). Because Rend-seq relies on cDNA circularization rather than RNA ligation to capture RNA 5′ ends, it is agnostic to the 5′ moiety generated following cleavage (Lalanne et al., 2018). To further confirm that these cleavage sites are driven by EndoA, which is encoded by the *ndoA* gene, we repeated these measurements upon deletion of *ndoA* in an RNase J1 deficient strain and found complete ablation of the cleavage signature in Rend-seq (Figure 4C, D). Furthermore, consistent with recent work indicating that EndoA recognizes a 6-mer motif and not a previously described 5-mer motif (Guler et al., 2024), we found that only U/ACAU followed by an adenosine leads to cleavage (Figure 4A, B).

**Figure 4.**
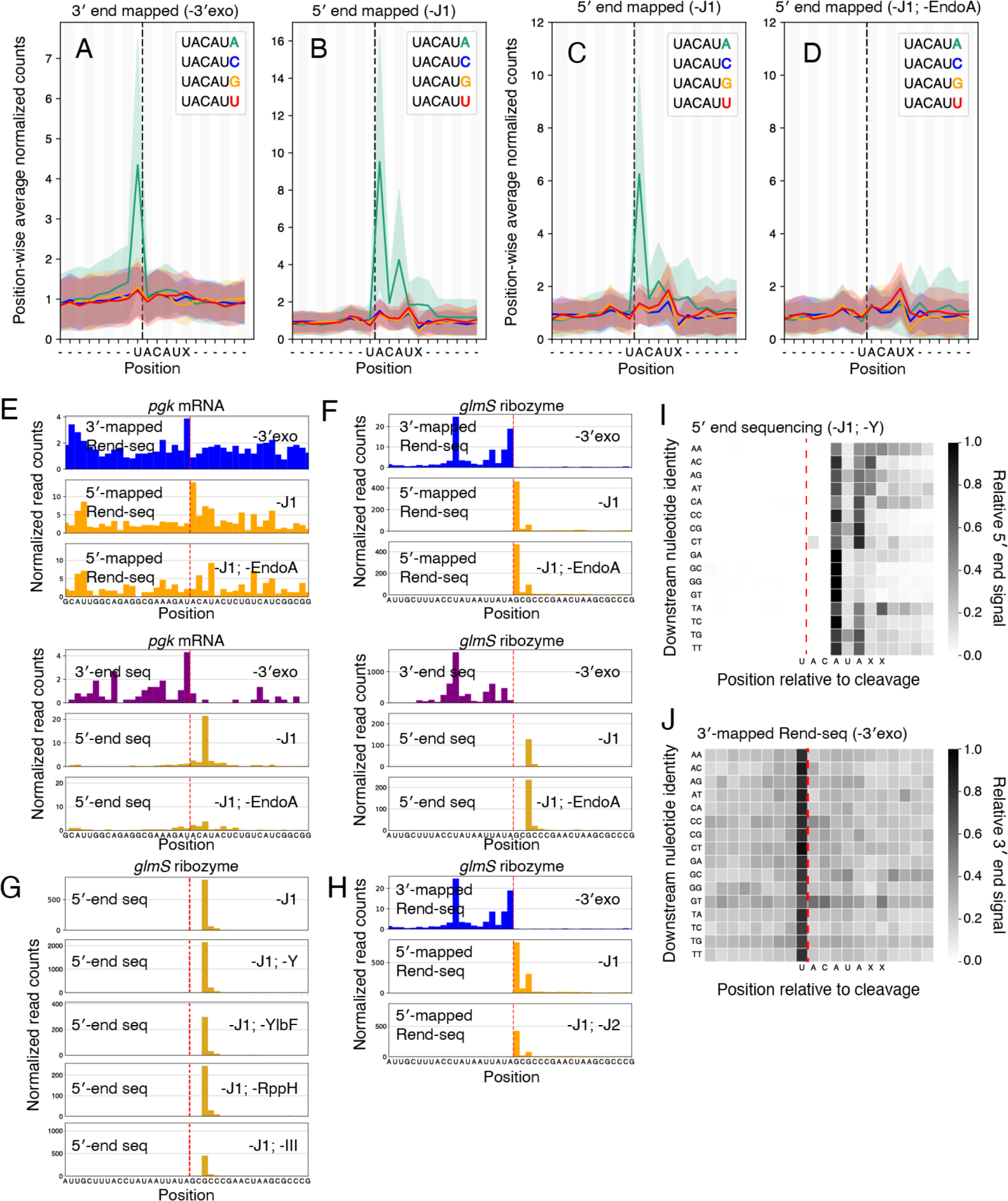
Revision of EndoA cleavage specificity and evidence for new ribonuclease activity in *B. subtilis*. (A, B) 3′ (A) and 5′ (B) mapped Rend-seq signal across all UACAU motifs in the genome, separated by downstream nucleotide. 5′ end sequencing data derived from an RNase J1 deletion (CCB434) and 3′ end sequencing derived from a 4-exo knockout (CCB396). Rend-seq data at each site is normalized to first 8 (for 3′-mapped) or last 8 (for 5′-mapped) positions within the window and a position-wise mean and standard deviation (shaded interval) are calculated with 90% winsorization. Motif instances with fewer than 1 read per position or fewer than 10 reads within the normalization window are not considered. Following this filtering, the number of considered sites ranges from 264 to 435 (A) or 292 to 455 (B) per motif. The black vertical line indicates the position of cleavage by EndoA. (C, D) 5′-mapped Rend-seq signal in an RNase J1 depletion (CCB390) (C) or and RNase J1 depletion with deletion of *ndoA* (BT231) (D). Analysis performed identically to (A, B). Following filtering, the number of considered sites ranges from 191 to 288 (C) or 86 to 117 (D) per motif. (E) Rend-seq (top) and end sequencing (bottom) data at a representative EndoA cleavage site located in the gene *pgk* for a 4-exo knockout (CCB396), RNase J1 depletion (CCB390), and RNase J1 depletion with an *ndoA* knockout (BJT231). 5′-mapped data shown in yellow (5′ end sequencing) or orange (Rend-seq) and 3′-mapped data shown in purple (3′ end sequencing) or blue (Rend-seq). Red dotted line indicates position of cleavage. Plotted are reads per million CDS-mapping reads. 5′/3′-end sequencing data are normalized to the average 3′-mapped Rend-seq signal in this window. (F) Rend-seq (top) and end sequencing (bottom) data at the position of cleavage by the *glmS* ribozyme. 5′-mapped data shown in yellow (5′ end sequencing) or orange (Rend-seq) and 3′-mapped data shown in purple (3′ end sequencing) or blue (Rend-seq). Strains included are identical to panel E. Red dotted line indicates position of cleavage. Data are plotted as in (E). (G) 5′-end sequencing data at the position of cleavage by the *glmS* ribozyme for knockouts of additional RNA decay-associated proteins. All knockouts are coupled to either a depletion or deletion of RNase J1. Knocked out genes include *rnjA* (RNase J1, strain CCB434), *rny* (RNase Y, strain CCB760), *ylbF* (YlbF, strain BJT074), *rppH* (RppH, strain BJT129), and *rnc* (RNase III, strain BG879). Data are plotted as in (E), with the exception of *rppH*, which is normalized to an *rnjA* knockout alone, rather than the same genetic background, due to a lack of corresponding Rend-seq data. (H) Rend-seq data at the position of cleavage by the *glmS* ribozyme in a 4-exo knockout (CCB396), RNase J1 knockout (CCB434), and RNase J1 depletion with a *rnjB* knockout (GLB186). Red dotted line indicates position of cleavage. 3′-mapped data are shown in blue and 5′-mapped data are shown in orange. Plotted are reads per million CDS-mapping reads. (I, J) 5′ end sequencing (I) and Rend-seq 3′-mapped (J) signal across all UACAUA EndoA cleavage motifs, separated by downstream sequence. 5′ end sequencing data derived from an RNase J1 depletion with *rny* knockout (CCB760) and 3′ end sequencing derived from a 4-exo knockout (CCB396). The signal in a 20 nt window around each cleavage site was normalized to its maximal value and a position-wise average across all normalized windows was calculated with 90% winsorization. UACAUA instances for which the local density in corresponding Rend-seq data is less than 1 read per position were discarded. Following this filtering, the number of considered sites ranges from 10 to 54 per row (I) or 10 to 62 per row (J). The red vertical line indicates the position of cleavage by EndoA.

Though our monophosphate-specific 5′ end sequencing approach did not detect the hydroxylated 5′ ends generated by EndoA, it unexpectedly captured peaks 2 nucleotides downstream of the canonical EndoA cleavage positions (Figure 4E). These 5′ monophosphorylated ends only appear when EndoA is present (Figure 4D, E), suggesting that two nucleotides are removed following cleavage by EndoA. To test if this trimming activity is general against all 5′ hydroxyl groups, not simply EndoA sites, we looked at the 5′ OH generated by the self-cleaving *glmS* ribozyme, whose downstream cleavage product is subject to subsequent degradation by RNase J1 (Collins et al., 2007; Winkler et al., 2004). In both a knockout of RNase J1 and a double knockout of RNase J1 and EndoA, 5′ end sequencing shows a clear 5′ monophosphorylated peak 2 nucleotides downstream of the site of self-cleavage in *glmS*, suggesting that this novel trimming activity is independent of EndoA (Figure 4F). The trimming activity is also independent of RNases Y, III, J1, the Y-complex constituent YlbF, or the RNA pyrophosphohydrolase RppH, as downstream peaks are readily detectable in strains lacking these enzymes (Figure 4G). Finally, this activity cannot be explained by RNase J2, as 5′-mapped Rend-seq measurements of a double knockout of RNase J1 and J2 also show a clear peak two nucleotides downstream of the canonical *glmS* cleavage position (Figure 4H).

We observed multiple rounds of 2-nt trimming at some, but not all, EndoA cleavage sites, suggesting sequence-specific trimming activities (Figure S4B,C). To determine the sequence specificity, we examined the downstream nucleotide content following the U/ACAUA EndoA motif (Figure 4I), focusing on our *rny* knockout and RNase J1 depletion strain. This strain was chosen for its particularly clear EndoA activity, which is likely driven by further stabilization of EndoA cleavage products following inactivation of an additional downstream decay pathway, though EndoA activity may also be increased. For sites with an EndoA motif followed by a U or C, monophosphorylated 5′ ends are found at both 2 and 4 nucleotides downstream of the initial cleavage, indicating discrete trimming (Figure 4I). For U/ACAUA motifs followed by an A (particularly by AA), additional 5′ ends are also generated five or more nucleotides downstream of the initial cleavage. By contrast, motifs followed by a G only produce 5′ ends at 2 nucleotides downstream. These 5′ ends are not driven by changes to the EndoA cleavage position, as the 3′ end density in the 4-exo knockout remains constant regardless of the downstream sequence context (Figure 4J). These data are consistent with successive rounds of nucleotide removal from the RNA 5′ end following endoribonucleolytic cleavage, with a preference for trimming sequences rich in adenosine and against those rich in guanosine. We are unable to rule out that differential stability following EndoA cleavage and 5′ end trimming may also influence the apparent sequence preferences we observe, however, and further work is required to reveal the identity and mode of action of this enzyme.

Cleavage 2 or 4 nucleotides downstream of the canonical EndoA cleavage site coincides with the U/A and C/A dinucleotides that are recognized by the common laboratory contaminant RNase A. Several lines of evidence argue against such a contaminant generating the trimming behavior we characterize in this work. First, RNase A cleavage generates RNA termini with 5′ hydroxyl or 3′ phosphate groups, similar to EndoA (Cuchillo et al., 2011). These ends cannot be captured by our RNA 5′ and 3′ end sequencing approaches (Figure 4E). Second, the 5′ trimming activity observed at *glmS* occurs within a C/G dinucleotide, suggesting that this activity is not restricted to U/A and C/A dinucleotides. Finally, our observation of 5′ end trimming is highly consistent across many replicate measurements of strains lacking RNase J1, as is an observed lack of trimming when this enzyme is present. It is thus highly unlikely that this trimming is driven by *in vitro* RNase activity downstream of cell lysis and RNA extraction.

### Determinants of cleavage by RNase Y

Though RNase Y is thought to be the primary endonuclease driving RNA decay in *B. subtilis*, only a small number of cleavage positions have been mapped in this bacterium, and what specifies RNase Y cleavage is poorly understood (Trinquier et al., 2020). To define the substrate features driving RNase Y cleavage, we sought to identify a set of putative cleavage sites that are dependent on the presence of this enzyme. For the 1330 putative endonuclease cleavage positions identified in our W168 background, we calculated their RNase Y sensitivity using RNase J1 deficient cells with or without an RNase Y knockout. This metric was bimodally distributed, and any cleavage signature with a sensitivity value over 10.56 was called as RNase Y dependent based on a double-Gaussian fit of these data (Figure 5A). Using this approach, we identified 596 cleavage sites across 387 genes that can confidently assigned to RNase Y sites (Table S4).

**Figure 5.**
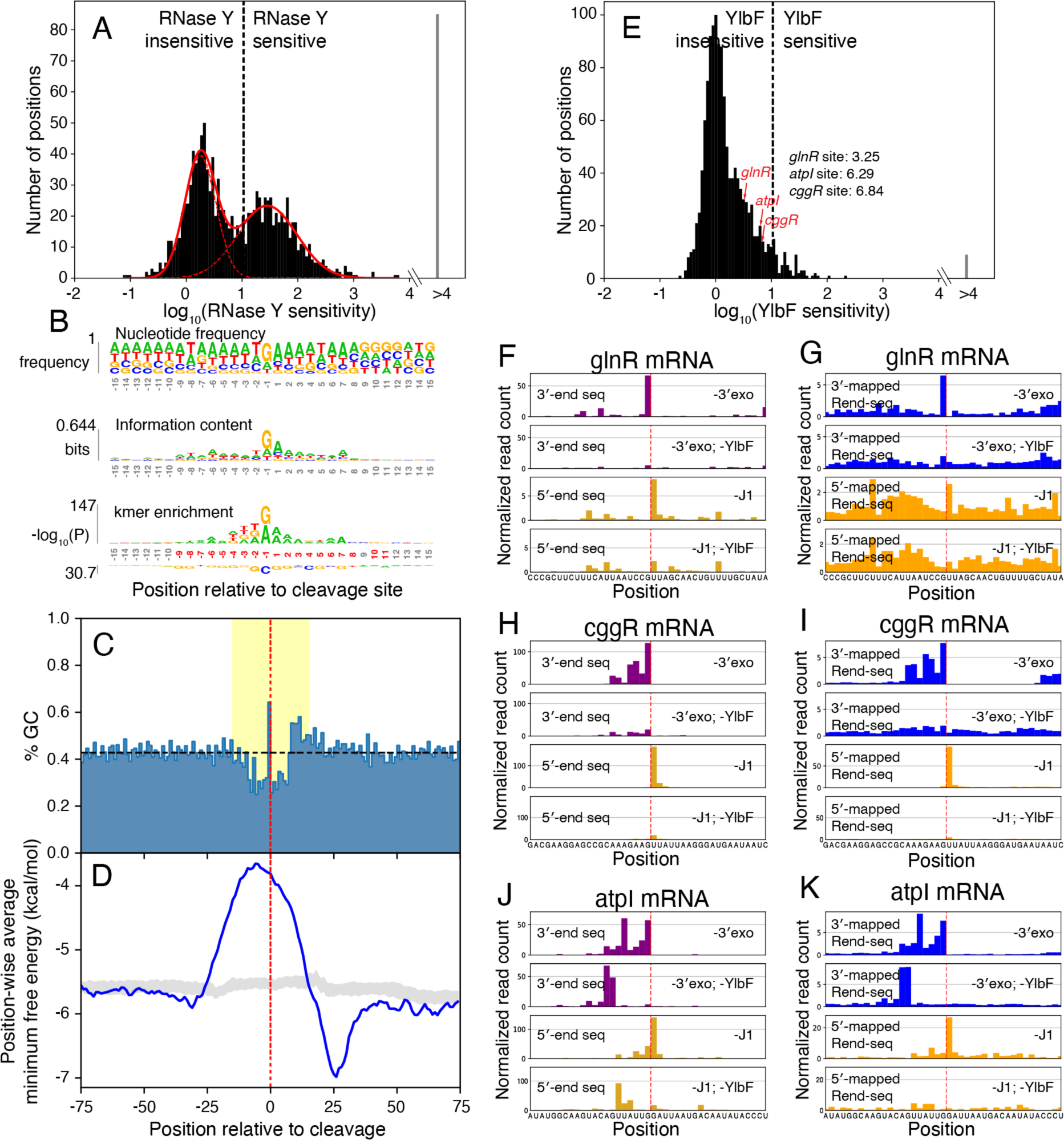
Sequence and structural features of putative RNase Y substrates. (A) Systematic identification of putative RNase Y cleavage sites. Histogram shows distribution of endonuclease sensitivities for called peak pairs in 3′/5′ end sequencing of exoribonuclease knockouts (see Figure S3 and Methods). Solid red line shows results of fitting the sum of two Gaussian distributions to this underlying log-transformed data, with the dashed lines showing the two constituent distributions produced by this fitting. The dashed vertical line indicates the lowest endonuclease sensitivity value (10.56) for which 95% of the Gaussian sum is derived from the right constituent distribution, a value above which sites are considered dependent on RNase Y. 596 of 1301 considered putative cleavage signatures exceed this threshold. For 85 sites we were unable to calculate a sensitivity score due to an absence of 5′ end sequencing counts in the knockout. These sites are called as RNase Y sensitive and are counted within the “>4” bin of the histogram. (B) Local sequence context around positions of RNase Y cleavage (N=596). Nucleotide frequency (top), information content (middle) and k-mer (bottom) logos in a 30 nucleotide window around called RNase Y sites. In k-mer logo, the most enriched k-mers are shown above the sequence and the most depleted below. Significant positions (Bonferroni corrected P<0.01, one-sided binomial test), are highlighted in red. (C) Local GC content around positions of RNase Y cleavage (N=484). Sites near transcript boundaries are not considered (see Methods). Dotted gray line indicates average %GC across *B. subtilis* genome. Vertical red line indicates position of cleavage. Yellow region indicates positions shown in (B). (D) Predicted folding near RNase Y cleavage positions (N=484). Sites near transcript boundaries are not considered (see Methods). Blue line shows position-wise average minimal free energy (MFE) predicted by RNAfold for 40-mers tiling cleavage sites. Gray band indicates per-position interquartile range of curves resulting from folding randomly sampled GA-dinucleotide centered windows within coding regions, excluding putative cleavage positions (see Methods). Red vertical line indicates putative position of cleavage. (E) Systematic identification of YlbF-dependent endoribonuclease cleavage sites. Histogram shows distribution of endonuclease sensitivities for called peak pairs in 3′/5′ end sequencing of exoribonuclease knockouts (see Figure S3 and Methods). The dashed vertical line indicates the endonuclease sensitivity threshold used to call RNase Y sites. Red arrows indicate endonuclease sensitivity values of canonical processing sites within the *cggR*, *glnR*, and *atpI* transcripts. For 9 sites we were unable to calculate a sensitivity score due to an absence of 5′ end sequencing counts in the knockout. These sites are called as YlbF sensitive and are counted within the “>4” bin of the histogram. (F-K) End mapping data showing evidence for inefficient but detectable endoribonucleolytic cleavage at canonical RNase Y processing sites within the transcripts encoding *glnR* (F, G), *cggR* (H, I), and *atpI* (J, K) in the absence of YlbF. (F, H, and J) show 5′/3′-end sequencing and (G, I, and K) show Rend-seq. 5′-mapped data shown in yellow (5′ end sequencing) or orange (Rend-seq) and 3′-mapped data shown in purple (3′ end sequencing) or blue (Rend-seq). Red dotted line indicates positions of endoribonucleolytic cleavage. Plotted are reads per million CDS-mapping reads. 5′/3′-end sequencing data are normalized to the average 3′-mapped Rend-seq signal in this window.

This set of putative RNase Y sites sheds light onto the substrate features that guide RNase Y activity. Frequency and information content logos constructed from the RNA sequences surrounding putative RNase Y sites suggest a sequence preference for cleavage downstream of a purine, with a particular preference for guanosine, and upstream of an adenosine (Figure 5B, top, middle). Similarly, considering k-mers of length ≤4, there is most significant enrichment for the dinucleotide sequence GA spanning the cleavage site, with an additional preference for uridine in the -2 and -3 positions (Figure 5B, bottom). This observed specificity is unlikely to be a result of biases in our methodology; identification of these cleavage sites requires concordance between 3′ and 5′ end sequencing datasets that are generated through distinct enzymatic reactions and adapter sequences. This sequence preference is additionally consistent with the sequence context in several previously mapped cleavage sites (Braun et al., 2017; Shahbabian et al., 2009), as well as the measured sequence preferences of RNase Y in *S. aureus* and *S. pyogenes*, despite the more limited role this enzyme plays in those organisms (Broglia et al., 2020; Khemici et al., 2015). However, the GA dinucleotide motif does not appear to be absolutely required, as 63% of the putative cleavage sites do not have this motif.

In a broader window around cleavage sites, there is a clear depletion for GC-containing sequence (Figure 5C) and a lower propensity to form secondary structures (Figure 5D). Computationally predicted folding of 40-nt RNAs surrounding positions of cleavage shows weaker structures for RNase Y sites compared to randomly sampled positions at uncleaved GA dinucleotides (Figure 5D). Interestingly, sequences centered just downstream of this lowly-structured region appear to fold more strongly than background, suggesting an enrichment for secondary structure proximal to the position of cleavage (Figure 5D). The downstream signal is recapitulated in the GC content within this window (Figure 5C), and is consistent with the influence of secondary structure on previously detected RNase Y cleavage events (Marincola and Wolz, 2017; Shahbabian et al., 2009). Taken together, these data indicate that *B. subtilis* RNase Y targets share a weak motif comprising a GA dinucleotide surrounded by AU-rich sequences that have reduced secondary structure.

### The complete Y-complex is not a strict requirement for RNase Y activity

Recent work has shown that the Y-complex, comprising proteins YlbF, YaaT, and YmcA, can act as a specificity factor for RNase Y for the maturation of nearly two-dozen operon mRNAs (DeLoughery et al., 2018). To understand if this complex plays a role in specifying the observed RNase Y cleavage events, we deleted the *ylbF* gene in both an RNase J1 knockout (CCB434) and 4-exo deficient background (CCB396). We then repeated our Rend-seq and end-sequencing measurements in these backgrounds, calculating the sensitivity of each putative cleavage site to YlbF (Figure 5E). Compared to RNase Y, few sites appeared dependent on the presence of YlbF, suggesting that the Y-complex plays a limited role in influencing RNA turnover outside of its known role in mRNA processing. Consistent with this observation, knocking out *ylbF* has a limited effect on the *B. subtilis* transcriptome (Figure S4D), as has been previously observed (DeLoughery et al., 2018).

Surprisingly, canonical maturation events, such as those within the *cggR-gapA*, *glnRA*, and *atp* operons are still observed in the absence of YlbF, albeit strongly attenuated (Figure 5E-H). Previously, the YlbF-dependence of these sites were determined using Rend-seq for cells that maintain the entire repertoire of exonucleases, which is less sensitive at detecting cleavage events. Here, by ablating exonucleases in cells lacking YlbF, we observed weak but detectable signals of cleavage products using Rend-seq, as well as unambiguous peaks in both 5′ and 3′ end sequencing (Figure 5F-K). These results underscore the sensitivity of our approach and suggest a revision of the role of the Y-complex. That is, instead of being a strictly necessary specificity factor for RNase Y cleavage, the Y-complex plays an enhancing role at these operon maturation events.

### Deep mutational scanning of RNase Y substrates

Our census of RNase Y cleavage events showed a degenerate motif, which is insufficient to explain its highly precise site selection for previously established mRNA targets. To identify additional sequence or structural features required, we developed a massively parallel reporter assay (MPRA) that allows high-throughput measurement of cleavage efficiency among a library of 10^4^-10^5^ mutated cleavage sites (Figure 6, Table S7). The reporter consists of a >100-nt region surrounding a cleavage site inserted into a stable native mRNA encoding the gene *aprE*, truncated such that it lacks an active site (Figure 6A). These reporter constructs are integrated at a neutral genomic locus (*ganA*) and contain a randomized 15-nucleotide barcode associated with each mutant variant, allowing relative quantification of each variant’s expression by RNA and DNA sequencing.

**Figure 6.**
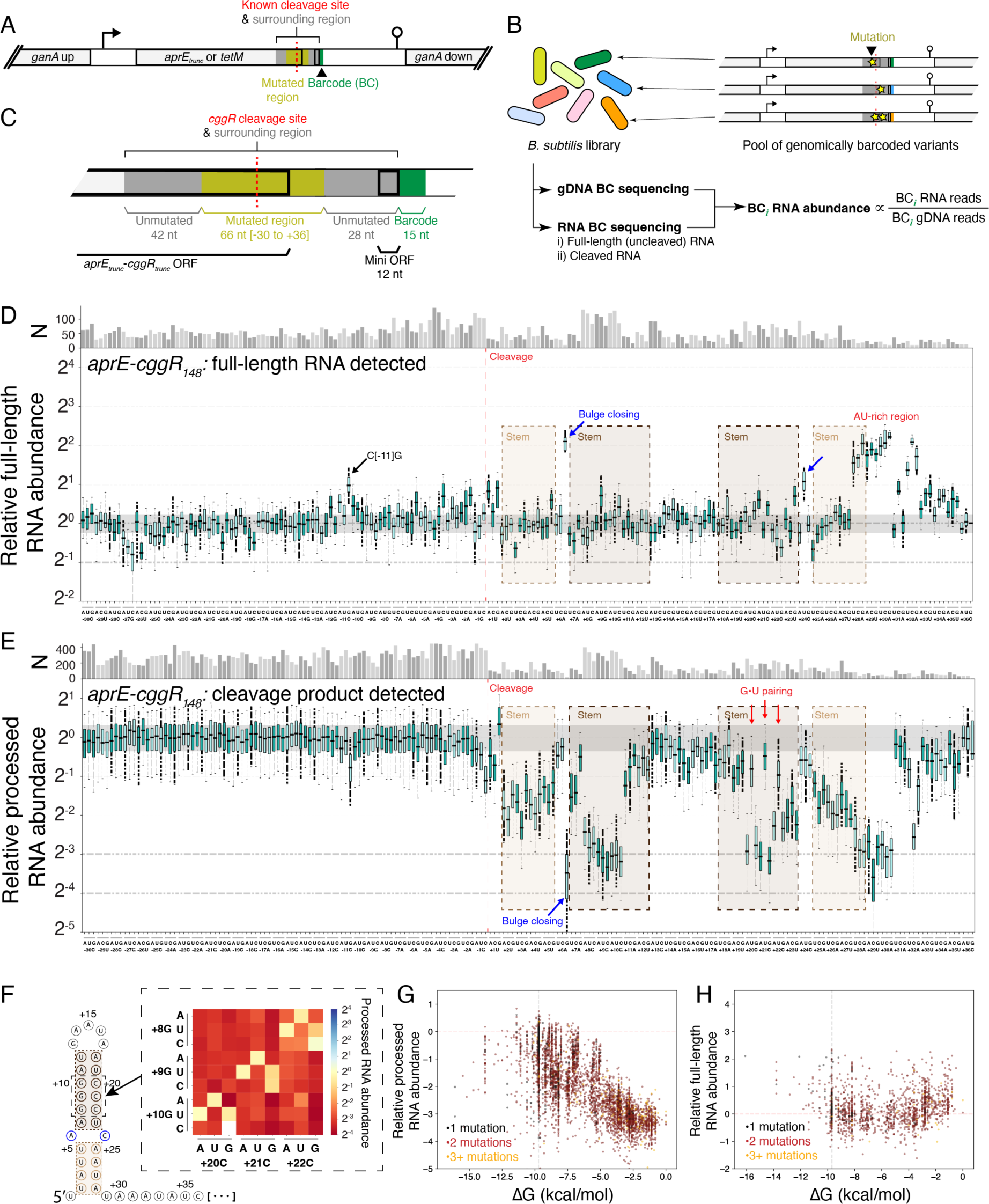
Mutational scanning of the *cggR-gapA* operon cleavage site. (A) Schematic design of MPRA construct. Cleavage region to mutate is indicated in dark gray with yellow used to indicate mutated region. Variant barcode indicated in green. Translated regions are indicated with a thick border. Promoters and transcriptional terminators are indicated by bent arrows and lollipops, respectively. (B) Workflow for measurement of mRNA processing. Protocol begins from *B. subtilis* culture containing a pool of genetically encoded cleavage site variants, and splits into gDNA and RNA barcode quantification protocols. The relative barcoded RNA abundance is calculated as the ratio of RNA-derived to gDNA-derived reads. (C) Schematic of the *cggR*_148_ construct inserted into the *aprE* MPRA transcript. *cggR*-derived sequence is colored gray, with the mutated positions highlighted in yellow. Translated regions are indicated with a thick border, and the variant barcode is indicated in green. (D) Impact of all single-nucleotide mutations on the accumulation of barcoded full-length RNA. Boxplots show variation between barcodes of identical variant sequence. Number of barcodes captured for each mutation is indicated above plot. Whiskers indicate 5^th^ and 95^th^ percentile. Gray shaded region indicates interquartile range for variants of wild-type sequence. Red line indicates position of cleavage. Blue arrows highlight mutations which close the bulge in the *cggR* downstream hairpin. Four variants (no more than one per mutation) have a value of zero are thus not visualized. (E) Impact of all single-nucleotide mutations on the accumulation of barcoded cleavage product. Boxplots show variation between barcodes of identical variant sequence. Number of barcodes captured for each mutation (N) is indicated by the bar height (top). Whiskers indicate 5^th^ and 95^th^ percentile. Gray shaded region indicates interquartile range for variants of wild-type sequence. Red line indicates position of cleavage and brown bars indicate regions predicted to pair in the formation of a downstream stem-loop structure (as illustrated in F). Red arrows indicate mutations which are predicted to generate G•U wobble pairing, and the blue arrow highlights a mutation which corrects the bulge in the downstream stem-loop structure (B). Four variants (no more than one per mutation) have a value of zero are thus not visualized. (F) Predicted stem-loop structure at 5′ end of processed RNA. Inset shows impact of double mutants on accumulation of processed RNA. Double mutant averages calculated over any variants with the two indicated mutations. Any variants which contain additional mutations that on their own result in greater than 10% change in processed RNA accumulation are discarded. (G) Relationship between predicted strength of downstream secondary structure and accumulation of barcoded cleavage product. The experiment each data point is derived from is denoted with a triangle (experiment 2b, Table S3), circle (4a), or square (6). The vertical dashed line indicates the ΔG of the unmutated sequence. Five variants fall outside of the bounds of this plot. (H) Relationship between predicted strength of downstream secondary structure and accumulation of barcoded full-length RNA. All data derived from experiment 2a (Table S7). The vertical dashed line indicates the ΔG of the unmutated sequence. Five variants fall outside of the bounds of this plot.

**Figure 7.**
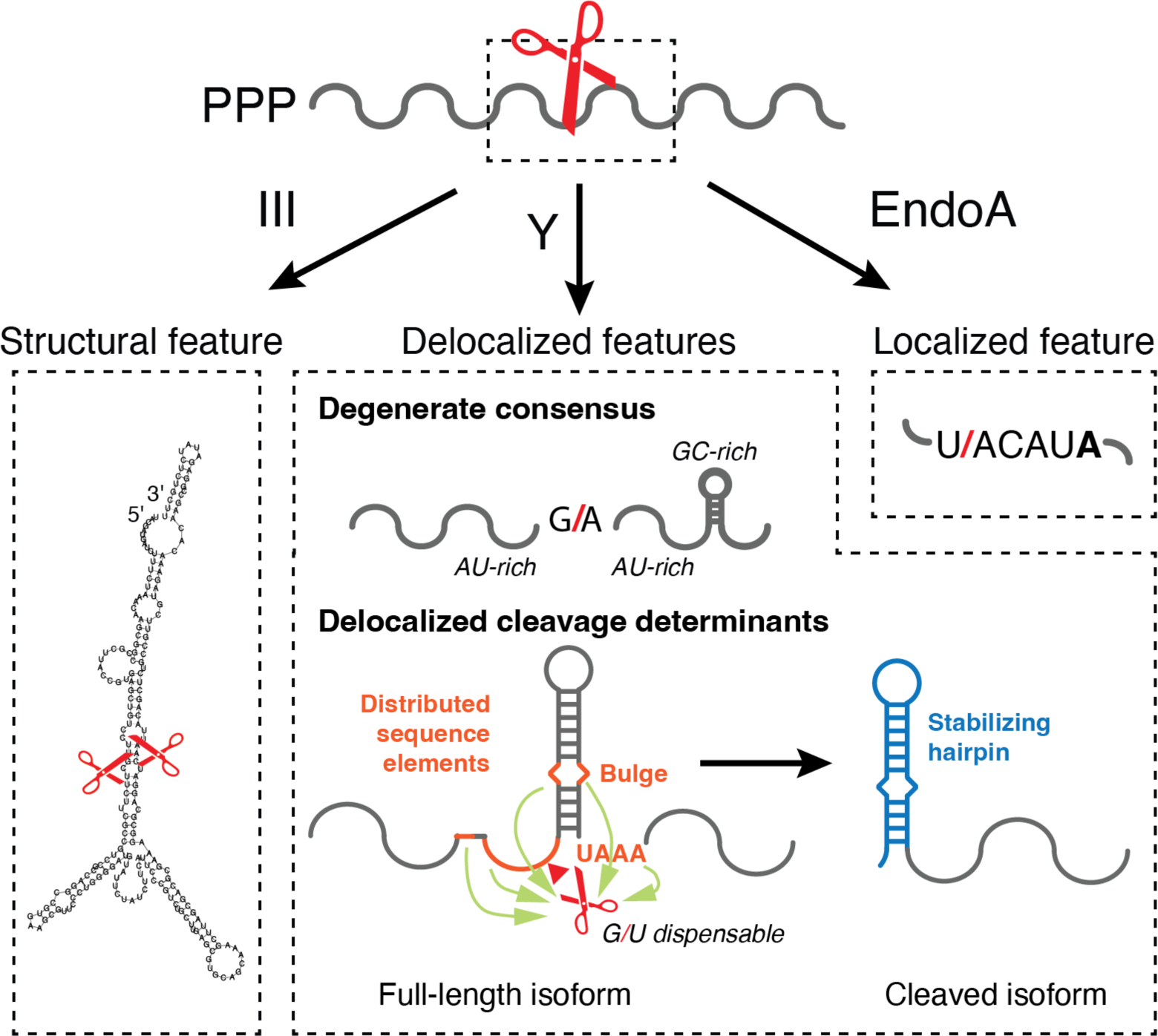
Determinants of endoribonuclease cleavage identified in this work. (A) Summary of the identified cleavage determinants of the three endoribonucleases profiled in this work. RNase III appears to cleave within double-stranded RNA sequences, including many duplexes formed through long range (>100 nt) intramolecular interactions, consistently leaving a 2 nucleotide 3′ overhang (left, “III”). EndoA appears to cleave specifically at a UACAUA primary sequence motif (right, “EndoA”). RNase Y cleavage appears to be targeted towards AU-rich regions of low secondary structure, appearing to prefer cleavage within a G/A dinucleotide (center, “Y”). Interrogation of the cleavage sites within *cggR* and *glnR* suggest that a -1G is dispensable for RNase Y cleavage, and for a given site the sequence and structural elements that drive cleavage (orange) may be distributed over many tens of nucleotides. Further, for some mRNA processing sites such as those in *cggR* or *gapA*, a downstream stem-loop structure (blue) appears to primarily drive stabilization of the cleaved isoform.

To simultaneously quantify cleavage activity at each mutant cleavage site, a pooled library of *B. subtilis* cells expressing our MPRA construct were grown and harvested in exponential phase for genomic DNA (gDNA) and RNA extraction (Figure 6B). Purified RNA is then subjected to reverse transcription and a size selection that separates cleaved from full-length RNA, allowing specific detection of each isoform. A reduction in cleavage should result in greater accumulation of the full-length RNA, with concomitant decrease in cleaved RNA. The barcode abundance in the uncleaved RNA fraction for each variant, normalized by its frequency in the pool as measured by gDNA sequencing, thus provides a readout of cleavage efficiency for that mutated sequence (Figure 6B).

### RNase Y cleavage is specified by distributed substrate features

We initially focused our MPRA measurements on a cleavage site found within the *cggR* gene, which in its native context drives differential expression within the glycolytic operon by producing a stable downstream product following cleavage (Ludwig et al., 2001; Meinken et al., 2003) (Figure 5F, G). The cleaved downstream product, which includes the *gapA* gene, is thought to be stabilized by an RNA hairpin structure at the cleaved end (henceforth referred to as the downstream hairpin) (Meinken et al., 2003), but whether proximal structure also plays a major role in specifying the cleavage site by *B. subtilis* RNase Y remains unknown. To capture surrounding sequences that may contribute to this cleavage, we inserted 148 bp of this sequence into our reporter, approximately centering the cleavage position within this window (Figure 6C). The site of cleavage, which in its native context resides in the *cggR* open reading frame, was similarly translated in the same reading frame of the reporter. Also adhering to the native context, the site of cleavage was followed by a stop codon and a short open reading frame derived from the start of the downstream *gapA* gene (Figure 6C). Using this approach, we characterized the effect of mutations from positions -30 to +36 relative to the cleavage position. After sequencing of the gDNA and RNA, 91,580 uniquely barcoded variants were detected across all experiments. These data cover virtually all possible single-nucleotide changes (Figure 6D) and 32% and 44% of all double-nucleotide changes across experiments detecting full-length and cleaved RNAs, respectively (Figure S5A, B). For each mutant, we average the effects over different barcode variants (median 44 barcodes for single-nucleotide changes) to reduce noise and barcode-driven variations (Figure S5C, D).

Mutations across a wide region surrounding the *cggR* cleavage site showed modest but detectable impact on steady-state levels of the full-length mRNA (Figure 6D). Between -30 and +1 nucleotide relative to the cleavage site, only a small number of mutations appear to reduce cleavage efficiency. The strongest effects are seen at the cytosine at position -11 (denoted as C[-11]), and the effect sizes are at most twofold. Interestingly, mutations of G at position -1 showed no statistically significant effect, even though it is the strongest consensus among RNase Y cleavage sites across the transcriptome. The fact that the G[-1] is not strictly required is consistent with our finding that 37% of the RNase Y cleavage sites detected do not have a guanine at this position (Figure 5B). Therefore, mutations upstream of the *cggR* cleavage site lend further support to our transcriptome-wide results that showed a weak consensus.

Downstream of the cleavage site, mutations in the hairpin region also produced limited effects on full-length RNA levels. Single-nucleotide substitutions in this region did not have significant effects (Figure 6D). Combinations of substitutions which further destabilize the hairpin also did not affect the apparent cleavage efficiency (Figure 6H). Surprisingly, the only mutations in this structured region that reduce cleavage are specific substitutions in a bulge of the downstream hairpin, including an A[+6]G substitution (fourfold) and a C[+24]U substitution (twofold) (Figure 6D). These two single-substitution mutants close the bulge and complete the hairpin, and other bulge-closing double substitutions at these two locations also showed reduced cleavage. Together, these results suggest that the downstream hairpin is not essential for cleavage, and that a stronger hairpin can in fact negatively impact cleavage.

Immediately downstream of the hairpin, an AU-rich region appears to play a role in cleavage. Mutations to AUAAAUAU located at position +28 to +35 lead to increases in full-length RNA level by up to 4-fold. In the folded RNA, this distal AU-rich region is brought to be adjacent to the cleavage site by the intervening hairpin (Figure 6F). Although it is plausible that such spatial proximity enables the distal region to recruit RNase Y or facilitate its activity, the fact that mutations in the hairpin region have no detectable effects suggests that other mechanisms may be at play.

To further elucidate the role of the downstream hairpin, we tested the hypothesis that this secondary structure mainly serves as a stabilizer for the cleaved product that encodes *gapA,* by preventing exoribonucleolytic decay (Meinken et al., 2003). This role is distinct from the model for the *Staphylococcus aureus cggR-gapA* operon, where the downstream hairpin has been suggested to specify RNase Y cleavage sites (Marincola and Wolz, 2017; Le Scornet et al., 2024). To distinguish these roles, we conducted deep mutational scanning on the accumulation of processed mRNA, measuring 349,266 mutant variants in total, and compared it to the results on full-length mRNA levels. Unlike full-length mRNA, the level of the downstream processed mRNA should depend on both cleavage efficiency and its own half-life post-cleavage.

Indeed, most mutations observed to increase full-length mRNA level resulted in reduced accumulation of the cleavage product, but not vice versa (Figure 6E). The effects of C[-11] mutations, bulge-closing mutation, and distal AU-rich region are recapitulated when we detected the downstream cleavage product. However, in contrast to the full-length RNA, mutations in the downstream hairpin region were among the most deleterious to cleavage product accumulation. Consistent with the secondary structure playing a role, double-substitution variants that complement these mutations (Figure 6F) have dramatically reduced effect sizes, and so are stem mutations that maintain base pairing (through wobble interactions, Figure 6E, red arrows). Considering all variants, including those with two or more substitutions, we found a clear trend that a stronger downstream hairpin is predictive of the cleaved RNA level (Figure 6G), in stark contrast with a lack of correlation for full-length RNA (Figure 6H). Finally, we repeated our measurements of cleaved RNA abundance in the context of an RNase J1 depletion, and found that, with the exception of closing the bulge, substitutions within the downstream stem no longer have a measurable impact when exoribonucleolytic activity against the processed RNA is attenuated (Figure S6A). Together, these results suggest that the downstream hairpin plays a much more substantial role dictating cleavage product abundance than the efficiency of RNase Y cleavage.

To assess whether the observed role of proximal secondary structure is specific to the *cggR* operon, we repeated the described measurements and analysis for a second cleavage site, located within the *glnR* gene of the *glnRA* operon (Figure S7A). As before, this cleavage occurs after the first gene in an operon and a stem-loop structure is predicted to form at the 5′ end of a stabilized downstream cleavage product (Figure S7B). Measuring accumulation of the cleaved RNA, we captured 54,431 variants with mutations occurring from positions -5 to +31 relative to the position of cleavage. For nucleotides inside the stem, both single mutations (Figure S7C) and higher-order substitutions that disrupt this RNA folding (Figure S7D) attenuated the accumulation of the cleavage product (up to approximately 4-fold), recapitulating the pattern seen for *cggR*. When we quantified the accumulation of full-length RNA (Figure S7E) or cleaved RNA upon RNase J1 depletion (Figure S7F), we found that disruption of the stem did not appear to stabilize the full-length RNA. Together, these results suggest that the behavior seen for *cggR* may be shared with other similarly structured mRNA processing sites.

Finally, we placed the *cggR* cleavage site into a second reporter construct encoding the exogenous gene *tetM* to assess whether the features driving cleavage product accumulation are context-dependent. Although most key effects are observed in both reporter constructs, the *tetM* reporter showed deleterious substitutions in C[-19]G and, to a lesser degree, A[-6]G (Figure S6B). The nucleotide C[-19] is situated in an otherwise purine-rich region, AGA **C** GAAGGAG, inside the *cggR* open reading frame. We initially hypothesized that the C[-19]G substitution may affect cleavage by slowing down elongating ribosomes through its Shine-Dalgarno-like sequence. However, the deleterious effects were still observed when we introduced an upstream stop codon, suggesting that these substitutions do not act through translation but other context-specific means (Figure S6C). These results reinforce the notion that the determinants of RNase Y cleavage are not localized to a narrow region on the mRNA.

## DISCUSSION

In this work, we present the first genome-scale map of *B. subtilis* endoribonuclease cleavages at single-nucleotide resolution. Using stabilized end-sequencing, we identified over a thousand putative cleavage events, allowing us to learn substrate preferences of the central endonucleases involved in *B. subtilis* mRNA decay. This work revealed the importance of long-range RNA base pairing for RNase III cleavage and confirmed that the toxin EndoA recognizes a hexameric rather than pentameric motif *in vivo*. We additionally uncovered evidence for and potential specificity of a new ribonucleolytic activity previously unknown in *B. subtilis*, motivating further work to understand RNA 5′ end processing and decay. Finally, we identified hundreds of new positions of cleavage by the core endonuclease driving mRNA decay, RNase Y. Combining this global cleavage mapping with a new massively parallel reporter assay, we demonstrate that cleavage by this enzyme is specified by features distributed over many positions across regions tens of nucleotides in length. This stands in stark contrast to enzymes like EndoA, which recognizes a strict consensus cleavage sequence, and RNase III, which has a strongly stereotyped cleavage behavior within secondary structures.

The broad suite of features that contribute to RNase Y processing at the *cggR* site helps explain the highly degenerate motif that we observed through global cleavage site mapping. Our data support a model in which RNase Y cleavage is localized not just by the sequence proximal to the cleaved bond, but rather by combined contributions of features distributed as many as tens of nucleotides away from the cleavage site. The ability for features distal in primary sequence to strongly influence cleavage may be mediated by RNA folding or by higher-order RNA-protein and protein-protein interactions between these features and components of the bacterial RNA degradosome. As such, the particular positions recognized to specify cleavage at each site may be variable and highly context-specific. Interestingly, a common feature we identified between many RNase Y sites, the presence of a -1G, only have a modest effect on the accumulation of processed *cggR* RNA in our reporter system, if any. This discrepancy may suggest a similar model to what is observed in in *S. aureus,* where an upstream guanosine fine-tunes the position of cleavage but is not required for RNase Y activity (Le Scornet et al., 2024).

The absence of a clear consensus motif specifying RNase Y activity echoes previous findings for RNase E in *E. coli* (Chao et al., 2017; Lin-Chao et al., 1994; McDowall et al., 1994). These observations suggest that the code for RNA decay fundamentally differs from transcription and translation, whose rates are influenced by well-defined interactions between the gene expression machinery and localized sequence features within either promoters or ribosome binding sites (Taggart et al., 2021). Instead, decay-initiating RNases such as RNase Y appear to be strongly influenced by features distal to the position of cleavage, possibly facilitated by RNA secondary structure and RNA-protein interactions. mRNA stability may also be strongly influenced by features extrinsic to primary RNA sequence. Decay can occur concurrently with and may be influenced by transcription and translation (Condon, 2003; Yarchuk et al., 1992), and the nonuniform distribution of decay machinery in bacterial cells also means that protein binding and RNA localization may have a strong influence on RNA stability (Hamouche et al., 2020; Lehnik-Habrink et al., 2011b; Moffitt et al., 2016). Robust prediction of endoribonuclease cleavage from primary sequence will thus likely require integration of more modes of measurement and interrogation of more cleavage sites than our current end-mapping approach and MPRA allow.

Although the presented global map of endonuclease activity expands our list of known endoribonuclease cleavage positions by orders of magnitude, we do not expect it to be comprehensive. First, due to size-selection steps in the end-sequencing protocol, we are unable to detect cleavage sites within 15 nucleotides of a 5′ end or 40 nucleotides of a 3′ end of a transcription unit. Additionally, while all known exoribonuclease activity in one direction is ablated in each of our experiments, secondary endoribonuclease cleavage followed by activity of the unperturbed exoribonucleases acting in the direction opposite may destabilize cleavage products, rendering them undetectable. Finally, activity by unknown exoribonucleases may cause broadening or shifting of 3′ or 5′ ends, separating the peak pairs and obscuring potential cleavage positions (as seen in Figure 4). The absence of signal in these data therefore is not necessarily an indication that turnover of a particular RNA is independent of endoribonucleolytic cleavage.

Unexpectedly, our analysis of EndoA cleavage sites, as well as the *glmS* ribozyme, revealed a previously uncharacterized 5′ end trimming activity in *B. subtilis.* We have confirmed that this activity is not dependent on the primary enzymes targeting mRNAs for decay, including RNases Y, III, J1, J2, RppH, or EndoA. Whether this nuclease acts in a 5′-3′ exonucleolytic manner or through endoribonucleolytic cleavage near 5′ ends remains to be determined. Further work is additionally necessary to understand whether this enzyme acts to facilitate RNase J1 cleavage at substrates with a 5′ hydroxyl moiety, as such behavior would shape our understanding of how systems like the *glmS* ribozyme regulate gene expression. Determining what gene encodes this activity is thus of great interest moving forward.

Our finding that the full Y-complex is not strictly required for processing of *B. subtilis* operons suggests a refinement of our understanding of this complex in RNA maturation and decay. Although our data support that loss of a component of the Y-complex strongly attenuates cleavage at the operons it canonically regulates, persistence of weak cleavage signatures suggests that the presence of YlbF is not a strict requirement for RNase Y activity at these sites. This residual cleavage may represent dispensability of the Y-complex as a whole or dispensability of YlbF itself for Y-complex activity. Indeed, recent work by Dubnau *et al*. has suggested that, of the Y-complex members, only YaaT (RicT) stably associates with RNase Y to form the functional complex required for operon maturation (Dubnau et al., 2023). It remains to be tested whether deletion of YaaT eliminates the low-level cleavage activity we detect in the absence of YlbF.

Mutational scanning of the *cggR* mRNA processing site revealed that downstream secondary structure primarily influences *cggR* operon stoichiometry through stabilization of the product of cleavage rather than driving cleavage itself. This result, while consistent with early studies of this operon in *B. subtilis* (Meinken et al., 2003), may appear to contradict studies in *S. aureus* that suggest presence of a downstream hairpin is a key driver of RNase Y cleavage (Marincola and Wolz, 2017; Le Scornet et al., 2024). It is worth noting, however, that *S. aureus* RNase Y targets a more limited set of RNAs than its *B. subtilis* homolog and regulation of cleavage may differ between the two enzymes (Khemici et al., 2015; Marincola et al., 2012). Additionally, the single base substitutions investigated here are a milder perturbation than used in other studies of the *cggR* operon (Le Scornet et al., 2024), and when secondary structure is strongly ablated in our experiments, a mild stabilization of the full-length construct is observed (Figure 6H). It is thus possible that although the effect of secondary structure on cleavage product stability is more pronounced, downstream RNA structure is playing a dual role in this system.

The approach described in this work has the potential to be generalized to study RNA decay in other bacteria for which exonuclease activity can be ablated or transiently depleted, representing a new approach to discover factors influencing RNA decay. Additionally, application of this approach to conditions beyond exponential growth in rich medium will provide an insight into regulation of endonuclease activity and RNA decay more broadly. The high-resolution picture of RNA decay initiation provided by this approach, particularly when integrated with future measurements of RNA half-life, will inform future quantitative predictions of RNA stability and gene expression levels from primary sequence in Gram-positive bacteria.

## MATERIALS AND METHODS

### Strains and strain construction

For all Rend-seq, 5′ end sequencing, and 3′ end sequencing experiments, excluding the Rend-seq measurement of an *rnjB* deletion with RNase J1 depletion, derivatives of W168 were used (Table S1). All transformations for strain construction were performed using standard protocols relying on natural *B. subtilis* competence and flanking homology mediated recombination using the media and protocol described in (Parker et al., 2020). BJT070 was created through SPP1 phage transduction of the *ylbF::erm* allele from BJT034 into CCB396 (Yasbin and Young, 1974). To create BJT074, gDNA from BJT034 was first transformed into SSB1002 to generate a *ylbF* knockout in a W168 background, BJT069. From this, SPP1 phage transduction from CCB434 was used to construct an *rnjA ylbF* double knockout, BJT074. CCB390 was constructed by first transferring an inducible RNase J1 allele (first described in (Britton et al., 2007)) into W168 (strain SSB1002), followed by deletion of *rnjA* through transformation with gDNA from CCB434. To construct BJT231, genomic DNA from GLB230 was transformed into CCB390, supplementing with 2% xylose in the liquid and solid media throughout the transformation protocol. To construct BJT129, CCB390 was transformed with genomic DNA from an *rppH* knockout, strain BKE30630 (Koo et al., 2017).

A supercompetent *B. subtilis* strain BJT200, based on (Zhang and Zhang, 2011), was used for experiments that do not require depletion or induction of RNase J1. This strain was constructed from W168 *B. subtilis* strain SSB1002 through insertion of an anhydrotetracycline (aTc) inducible *comK* allele at the *ganA* locus by transformation of ScaI-linearized plasmid pJT117. For experiments in which an inducible RNase J1 was required, strain BJT201 was utilized. Similar to BJT200, this strain contains an inducible *comK* gene. BJT201 is a derivative of strain CCB390, a W168 *B. subtilis* strain with xylose inducible RNase J1, constructed through transformation with pJT117. These strains were used for all MPRA experiments except the *tetM-cggR* data, which were collected in a *trpC* 168 strain. Plasmid pJT117 and MPRA plasmids pJT028 and pJT124 were constructed by isothermal assembly.

### Cell growth and harvesting (Rend-seq, 5′-, and 3′-end sequencing)

All Rend-seq, 5′-, and 3′-end sequencing experiments were performed using cells grown in 2xYT, supplemented with 2% xylose or glucose for RNase J1 induction or depletion as necessary. Except when we sought to deplete RNase J1, strains with an inducible RNase J1 allele were consistently grown in the presence of 2% xylose. For all sequencing experiments, overnight cultures in 2xYT were back-diluted to an OD_600_ of 0.0002 in 20 mL prewarmed 2xYT and grown to an OD_600_ of 0.2, at which point they were harvested. For any strain with an RNase J1 knockout, cultures were vortexed prior to and following inoculation to minimize clumping. To achieve RNase J1 depletion, 10 times the required volume of overnight culture was collected, washed twice with 1 mL pre-warmed 2xYT, and resuspended in 1 mL 2xYT. 100 µL of this resuspension was used to inoculate the final culture. Samples were harvested for sequencing by adding 7 mL of culture to 7 mL of ice cold methanol, pelleting, decanting, and flash freezing with liquid nitrogen. Cell pellets were stored at -80°C prior to RNA extraction.

### End-enriched RNA sequencing and 3′ end sequencing

Prior to RNA extraction, with the exception of our first Rend-seq experiment on the exonuclease-deficient strains (profiling wild-type 168, 4-exo, and Δ*rnjA* strains with and without *ylbF* knockout), cell pellets were washed with 10-15 mL ice cold 10 mM Tris pH 7.0 to remove contaminating nucleic acids that precipitate from 2xYT in methanol. For Rend-seq, RNA was extracted from cell pellets using either an RNeasy Mini Kit (Qiagen) with on-column DNase treatment, an RNeasy Plus Mini Kit (Qiagen) or through RNAsnap extraction (Stead et al., 2012) followed by Turbo DNase treatment. All 3′ end sequencing samples were purified through RNAsnap. In the RNAsnap protocol, cells were resuspended in 500 µL extraction solution (95% Formamide, 1% BME, 0.025% SDS, 18 mM EDTA) and transferred to a 2 mL tube containing 200 µL chilled 0.1 mm diameter zirconia beads. These tubes were vortexed for 10 minutes using a horizontal vortex adapter and subsequently incubated at 95 °C for 7 minutes. Samples were spun at 15,000 g, after which 400 µL was removed and diluted 1:5 with 1.6 mL DEPC-treated water. This mixture was precipitated with isopropanol and resuspended in 10 mM Tris, pH 7.0. At least 30 µg of RNA was then treated for 30 minutes at 37°C with 10 µL Turbo DNase in a 100 µL reaction. Following RNA extraction and gDNA depletion, 20 µg of purified RNA was treated with MICROBExpress Bacterial mRNA Enrichment Kit (ThermoFisher) per the manufacturer’s instructions and precipitated with isopropanol.

Rend-seq libraries were prepared as described previously (Lalanne et al., 2018). In short, rRNA depleted RNA was fragmented for 25 seconds (ThermoFisher #AM8740) and fragments of sizes 15-45 were size selected on a 15% TBE-Urea polyacrylamide gel (ThermoFisher). RNA was dephosphorylated with T4 PNK (NEB) and 3 pmol of this dephosphorylated RNA was ligated to an adenylated adaptor using truncated T4 RNA ligase 2 K277Q (Jonathan Weissman). Ligated RNA was purified by denaturing polyacrylamide size selection and reverse transcribed using SuperScript III (ThermoFisher) with a custom primer oCJ485. This cDNA was then circularized using CircLigase (Lucigen) and barcodes and full Illumina adapters were added through PCR using Phusion (NEB).

3′ end sequencing libraries were prepared in a manner similar to Rend-seq, with deviation in the order of reactions (Herzel et al., 2022). Prior to fragmentation, 1.8 µg of rRNA depleted RNA was ligated to an adenylated adapter as described for Rend-seq. This ligation product was then fragmented and RNA species of sizes 33-63 nt were purified via denaturing polyacrylamide gel extraction. From this point onwards, the library preparation is identical to Rend-seq. Note that this protocol lacks a dephosphorylation step prior to 3′ end ligation, such that RNA ends with a 3′ phosphate will not be captured by this protocol. Both Rend-seq and 5′ end sequencing libraries were sequenced with 50 or 75 bp single end reads on either a HiSeq 2000 or NextSeq500.

### Multiplexed 5′ end sequencing

For all 5′ end sequencing samples, pellets were washed, and RNA was extracted using the RNAsnap protocol and Turbo DNase treated as described for the Rend-seq and 3′ end sequencing experiments. rRNA was depleted from 10 µg of purified RNA with the MICROBExpress Bacterial mRNA Enrichment Kit, per manufacturer’s instructions, precipitated, and resuspended in 10 mM Tris pH 7.0. 600 ng rRNA depleted RNA was then ligated to 260 pmol of the RNA adapter oJT900 (Table S2) for 18 hours at 22 °C using T4 RNA Ligase 1 (based on (Subtelny et al., 2014)).

Following ligation, RNA was fragmented for 2 minutes (ThermoFisher) and fragments of either size 41-71 (corresponding to 15-45 plus a 26-nucleotide adapter) or 60-90 were isolated through denaturing polyacrylamide gel extraction. The larger size window was chosen to increase effective sequencing depth by removing dimerized fragments of oJT900, which were observed in high abundance in sequencing libraries prepared with a 41-71 nt size selection. RNA fragments were dephosphorylated, polyadenylated, and reverse transcribed following the BaM-seq multiplexed RNA sequencing protocol (Johnson et al., 2023). In short, in a single tube RNA fragments are sequentially dephosphorylated with T4 PNK (NEB), polyadenylated with *E. coli* poly(A) polymerase (NEB) and reverse transcribed with SuperScript III (ThermoFisher) and a barcoded reverse transcription primer. The cDNA from each sample generated by this reaction can then be pooled and purified by denaturing polyacrylamide gel electrophoresis. To add Illumina adapters, pooled cDNA was amplified between 5-10 cycles with primers oDP128 and oDP161 (Johnson et al., 2023) using Q5 DNA polymerase and purified by polyacrylamide gel electrophoresis. 5′ end sequencing libraries were sequenced with 75 bp single end reads on a NextSeq500.

### Rend-seq, 5′-, and 3′-end sequencing data processing

Adapter or poly(A) sequence was trimmed from end sequencing and Rend-seq libraries using CutAdapt (Martin, 2011) with parameters “-a ’CTGTAGGCACCATCAAT’ -m 15” or “-a A{15} -m 15 -u 8 -- rename=’{id} UMI={cut_prefix}’”. 5′ end sequencing reads were additionally collapsed using the 8 bp unique molecular identifier (UMI) at the start of each read using custom Python scripts, requiring identical UMI sequences and read sequences within a hamming distance of 1 to be considered duplicates. Trimmed reads were then mapped to the *S. cerevisiae* genome (SacCer3) using bowtie v1.1.2 with parameters “-v 1 -k 1 –best –un” with retained unmapped reads subsequently mapped to the *B. subtilis* genome (NC_000964.3) with the same parameters (Langmead et al., 2009). Mapping against yeast removes contaminating 2xYT-derived sequences and allows estimation of the degree of contamination in a particular sample. Using custom Python scripts, we converted the bowtie output to wig files, attributing signal from reads with a mismatch at their first position to the adjacent nucleotide to address mismatches due to non-templated addition during reverse transcription.

### Identification of putative endonuclease sites

To identify putative sites of endoribonucleolytic cleavage, we identified positions for which an adjacent 5′ and 3′ end could be detected in our end sequencing approaches. To do so, we first normalized each dataset to the total number of millions of CDS-mapped reads. At each position, we identified peaks in read density by calculating a ratio of 5′ end signal at that position to the 90% winsorized average in a window 50 nucleotides upstream (for 3′ ends) or downstream (for 5′ ends) of that position, excluding a 2-nucleotide gap on either side of the peak. For any adjacent positions (allowing a gap of 1 position) with a ratio over background greater than 7.5, these positions were grouped for the purposes of downstream 5′ end signal quantification. At each position with >10 reads per million CDS-mapped reads (RPM) in 5′ end sequencing, we detected paired 3′ ends by calculating an equivalent ratio centered around the position one nucleotide upstream in the 3′ end sequencing of the 4-exo deletion. Additionally, in the 50-nucleotide window downstream of each position, we calculated the local density of mapped 3′ ends of the RNase J1 deletion Rend-seq data to establish that we were considering positions only in genes expressed in this strain. Putative cleavage sites were called as positions for which the average local Rend-seq depth exceeded 0.05 RPM and both the 3′ and 5′ end sequencing signals had a ratio of 7.5 over background.

Because the sequencing methods used in this work are unable to capture short RNA fragments (<15 nt for Rend-seq and 3′ end sequencing and <15-34 nt for 5′ end sequencing, depending on the exact size selection criteria used), we did not consider positions within 40 nucleotides of an RNA 3′ end. To identify these ends, we called peaks in 3′-mapped RNase J1 Rend-seq data using a z-score based peak calling method. At each position, a z-score is calculated as the signal at that position minus the average signal in a window of 50 nucleotides on either side of the peak (excluding a gap of 2 nucleotides on either side of the peak), divided by the standard deviation within this window. Any position for which the calculated score exceeds 8 is flagged as a peak (i.e. an RNA 3′ end) in our Rend-seq data. Putative cleavage positions located up to 40 nucleotides upstream of one of these 3′ end positions are discarded. To count the number of putative cleavage positions across all our datasets, positions that were called in either the W168 (CCB434) or phage-cured W168 (BG880) background were considered. If two adjacent positions were called across these datasets, they were counted only a single position.

To attribute putative cleavage positions to specific endoribonucleases, we calculated the ratio of 5′ end sequencing signal at each position in an endonuclease knockout with RNase J1 deletion (or depletion) to the average Rend-seq signal of the same strain in a corresponding window 50 nucleotides on downstream of that position, excluding a 2-nt gap. To account for differences in sequencing depth between samples, this 5′ end sequencing and Rend-seq signal at each position was normalized to the total CDS-mapped reads in each sample. Additionally, signal from positions which were previously flagged as grouped in the 5′ end sequencing were summed, with the cleavage assigned upstream of the 5′-most position in the group. This ratio of 5′ end sequencing to Rend-seq signal is calculated for the corresponding RNase J1 deficient strain that expresses the considered endonuclease (e.g. a phage-cured RNase J1 deletion when considering RNase III). The ratio in the endoribonuclease expressing strain is then divided by the ratio in the endoribonuclease knockout strain to produce a “sensitivity” score. This approach is similar to that used to map RNase III sites in previous work (DiChiara et al., 2016).

Putative cleavage positions with a sensitivity score that exceeds 10.56 in a particular endoribonuclease knockout are called as cleavage sites for that enzyme. This threshold was empirically determined by fitting of the distribution of log-transformed sensitivity values for the RNase Y knockout to the sum of two Gaussian distributions and finding the point at which 95% of this fit is contributed by the underlying Gaussian with the larger mean (see Figure 5A). For this endonuclease assignment analysis, any position for which the local Rend-seq density in the endonuclease knockout strain (as calculated above) falls below 0.05 RPM is excluded to avoid spuriously calling positions in genes that are downregulated in this condition. When calculating the fraction of putative cleavage sites that are dependent on a particular protein, only those cleavage sites that were identified in an RNase J1 knockout of the same background were considered (e.g. only sites appearing in the phage-cured RNase J1 knockout strain BG880 were used when calculating the fraction of sites sensitive to RNase III knockout, which was generated in the same phage-cured W168 background, see Table S1).

### Determination of RNase III site duplex length, genomic distance, and structural distance

To identify pairs of RNase III cleavage positions that might represent intramolecular pairing, we first identified all RNase III-dependent cleavage positions within 1 kb of one another. The distance between sites in the primary sequence is the genomic distance as reported in Figure 3D. We then extracted the intervening RNA sequence plus a 50 nucleotide buffer on either side for each pair of positions and calculated its minimum free energy (MFE) structure using RNAfold from ViennaRNA v2.4.14 with default parameters (Lorenz et al., 2011). We then used this MFE structure and a custom Python script to calculate the length of the overhang generated (if any) by RNase III cleavage at this pair of sites. This length is reported as structural distance in Figure 3E. The reported duplex length for each structure was manually annotated, allowing for a maximum gap of two unpaired bases between base pairs.

### RNase III sequence-structure motif analysis

For analyses of sequence content within RNA secondary structures recognized by RNase III, we considered an expanded set of putative cleavage sites, using a threshold of >2 reads per million CDS-mapping reads in 5′ end sequencing (rather than the previous >10 RPM). Results of putative cleavage position identification and endonuclease sensitivity calculation using this threshold are summarized in Tables S5,6. From this set of putative cleavage sites, identified pairs of sites within 1 kb of one another and calculated a MFE secondary structure as described above. As before, we then used custom Python scripts to quantify the overhang length generated by the two RNase III cleavages and retained any sites which produced a 2-bp 3′ overhang.

Using the predicted MFE structures for these pairs of sites, we identified the base pair at each position within the structure relative to the two positions of cleavage. The position-wise frequency of each type of base-pairing interaction (A-U, U-A, G-C, C-G, U-G, or G-U) and fraction of sites for which that position was paired was then determined. In cases where an asymmetric bulge was observed, those bases were included as additional unpaired positions, rather than skipping over them in our indexing of base-pairs. To compare our data to those previously-published for *E. coli* (Altuvia et al., 2018), we calculated the difference between the frequency of each pairing type in our data and the equivalent observed at that position by Altuvia et al.

### Local secondary structure prediction for RNase Y sites

To determine folding propensity of sequences proximal to endoribonuclease cleavage sites, the *B. subtilis* genome was tiled with 40-mers and folded with RNAfold from the ViennaRNA v2.4.14 package with default parameters (Lorenz et al., 2011). The MFE values generated from this folding were assigned to the center position within each k-mer and used to generate wig files containing positional folding propensity. These values were extracted in 150 nucleotide windows centered around putative endoribonuclease cleavage positions and a position-wise mean was calculated. If a cleavage site was identified within 75 nucleotides of a 3′ or 5′ end present in wild-type W168 Rend-seq data (identified in strain SSB1002 by the previously described z-score based peak calling method), that cleavage site was not included in this analysis. To generate a background model that controls for sequence context in the - 1/+1 positions, this analysis was repeated 100 times sampling an equivalent number of random positions within coding regions that are centered on GA dinucleotides (the most enriched dinucleotide flanking putative RNase Y sites), excluding sites called as cleavage positions. From these predicted position-wise average folding curves, the 25^th^ and 75^th^ percentile at each position across all curves was used to generate the shaded region on Figure 5D.

### Information content and k-mer logo generation

To generate sequence logos, the local sequence in a 30-nucleotide window surrounding all putative cleavage sites for a given enzyme were identified and aggregated into a FASTA file. A background set of 30-mers evenly sampled every 100 nucleotides from *B. subtilis* coding regions (with sampling beginning after the start codon) was generated as a background set. kpLogo v1.1 (Wu and Bartel, 2017) was then used to generate information content and k-mer logos with parameters “-bgfile %s -startPos 16 -pc 0.01”, with %s pointing to our background set.

### MPRA plasmid library construction

Plasmid libraries were constructed through restriction cloning of randomized mRNA processing site sequences into suitable reporter plasmids (pJT028 or pJT124 for *tetM* or *aprE* experiments). Inserts for plasmid libraries were constructed through a two-step protocol using mutated oligonucleotides generated by IDT as a template. Oligonucleotides were designed with a 9% error rate (3% per non-native nucleotide) over the specified positions (see Table S7 for enumeration of experiments and their variable regions). In the first step, 10 fmol of oligo was PCR amplified for 12 cycles using Q5 DNA polymerase (NEB). To produce a *cggR*-derived insert with mutations downstream of the cleavage site (experiments 2a, 2b, 4a, 4b, and 6), oJT837 was amplified with oJT851/852. To prepare the *glnR* insert (experiments 8a, 8b, 9), oJT839 was amplified with oJT857/858. For the *cggR* insert with upstream mutations (experiments 1a, 1b, 3, 5, 7), instead of PCR amplification, oJT592 and oJT593 were mixed and extended through one cycle of denaturing, annealing, and extension with Q5. These products were diluted 1:10 into a second reaction in which they were amplified and barcoded through 4 cycles of PCR with Q5 using oJT548/594 and oJT855/859 for all *cggR* and *glnR* inserts respectively. Amplified, barcoded inserts were then cleaned up with a DNA Clean & Concentrator 5 column (Zymo Research).

To construct the plasmid pool, the amplified insert and 4 µg plasmid backbone were digested with SalI-HF and NheI-HF for 60 minutes at 37°C and purified with a DNA Clean & Concentrator 5 column. 400 ng cut plasmid and insert were mixed at an approximately 1:10 ratio and ligated through a 5-15 minute treatment with QuickLigase (NEB) at 25°C. This reaction was then cleaned up with a DNA Clean & Concentrator 5 column and transformed into electrocompetent *E. coli* (NEB 10-beta or NEB 5-alpha cells). Transformants were plated on LB with 100 µg/mL carbenicillin in 245 mm square bioassay dishes (Corning) and grown at 37°C overnight. Cells were then scraped off the plates and pelleted for ZymoPure II Maxiprep plasmid extraction (Zymo Research).

### Transformation and integration of MPRA library into B. subtilis genome

To introduce our plasmid libraries into *B. subtilis*, we utilized an inducible competence system mediated by an aTc-inducible *comK* allele introduced into BJT200 and BJT201, based on (Zhang and Zhang, 2011). A single colony was picked into LB (with 2% xylose for BJT201) and allowed to grow to an OD_600_ of 1.0 at 37°C with vigorous shaking. At this point, 10 ng/mL aTc was added to the culture to induce expression of ComK. Cells were shaken at 37°C for an additional 2 hours to allow induction of competence, at which point 100 µL per mL culture of plasmid pool linearized with either ScaI-HF or AfeI (NEB) was added to the culture. This restriction digest was prepared per the manufacturers protocol with 1 µg plasmid per 50 µL reaction. The cells were cultured with DNA for 1.5 hours, at which point the cultures were pelleted at room temperature, resuspended in residual media, and plated on 245 mm bioassay dishes containing 100 µg/mL spectinomycin or 5 µg/mL chloramphenicol. The *cggR-tetM* library was transformed using standard protocols relying on natural competence. After an overnight growth at 37°C, cells were scraped into LB with the appropriate antibiotic and approximately 500 million cells were back-diluted into 200 mL LB with antibiotic. This culture was grown for approximately 4-8 hours, and several 1 mL aliquots of culture were mixed with equal volume 40% glycerol and frozen at - 80°C to prepare stocks for future experiments.

### Cell growth and harvesting (MPRA)

For all MPRA experiments, cells were grown in LB medium with an inducer (20 ng/mL anhydrotetracycline or 2% xylose) when noted. For all experiments in an inducible RNase J1 background, 2% xylose was consistently used to maintain RNase J1 expression, which was substituted for 2% glucose when RNase J1 depletion was required. To collect samples, a single cryovial of a *B. subtilis* library was thawed, washed twice with 10 mL LB to remove glycerol, and grown overnight in 200 mL of LB at 37°C in a 2.8 L flask with vigorous shaking. These cultures were then back-diluted to an OD_600_ of 0.0003 in 200 mL LB with library inducer and grown at 37°C until an OD_600_ of 0.3, at which point samples were harvested. Two cell pellets were harvested per sample, one for gDNA extraction and one for RNA extraction. For RNA, 7 mL culture was added to 7 mL ice cold methanol, pelleted, decanted, and flash-frozen in liquid nitrogen. For gDNA, 14 mL of culture was pelleted, decanted, and frozen in liquid nitrogen. Cell pellets were stored at -80°C until nucleic acid extraction.

### MPRA RNA and DNA sequencing

Three sequencing libraries were prepared for each sample: one to quantify the barcodes in cellular RNA, one to quantify barcodes in the gDNA, and one to map barcodes to variant sequence. To prepare the RNA barcode quantification library, RNA was first extracted from frozen cell pellets with gDNA depletion using an RNeasy Plus Mini Kit (Qiagen). For experiments 1a, 1b, 2a, 2b, and 9 (see Table S7), 40 µg of total RNA was reverse transcribed using primer oJT442 and SuperScript III. This cDNA was then run on a 6% TBE-Urea gel, and the following size selection ranges were gel purified: 300-500 nt for full-length *cggR* and *glnR* constructs (experiments 1a, 2a, 9) and 100-250 nt for cleaved *cggR* constructs (experiments 1b and 2b). For the remaining experiments, cleaved RNA was isolated by size selection of RNAs ranging from 200-400 nt for *tetM-cggR* sites, 150-300 nt for *aprE-cggR* sites, and 150-350 nt for *aprE-glnR* sites on a 6% TBE-Urea gel (Novex). The size selected product was reverse transcribed using primer oJT442 and SuperScript III (Thermo Fisher) and size selected on 6% TBE-Urea gel, extracting approximately 120 to 200 nucleotides for *cggR* libraries and 100 to 200 for *glnR*. For all experiments, size-selected cDNA was amplified using Phusion DNA polymerase (NEB), adding a barcode and Illumina adapters with a Truseq indexing primer and an isoform-specific primer (see Table S2 for primer sequences). These isoform-specific primers were oJT441/oJT1290 for cleaved/full-length products from *cggR* constructs or oJT890/oJT1291 for cleaved/full-length products from *glnR* constructs. The resulting libraries were sequenced with 50 or 75 bp single end reads on an Illumina NextSeq500.

To prepare both libraries for gDNA barcode quantification and barcode to variant sequence mapping, a single gDNA sample was prepared from a frozen cell pellet using the Wizard Genomic DNA Purification Kit (Promega). Both gDNA-derived libraries were generated using a two-step Phusion PCR protocol. An initial amplicon was generated through 2 cycles of PCR using from 4 µg of purified nucleic acid per 50 µL PCR reaction (reaction scaled up as needed to minimize UMI collapse in more complex libraries) with primers oJT444/445 for *cggR* upstream mutations, oJT1125/445 for *cggR* downstream mutations, and oJT891/445 for *glnR* downstream mutations. The untranslated *cggR* library required a longer amplicon to capture the upstream stop codon and was instead amplified with oJT942 and oJT445. This first PCR reaction was cleaned up with a double-sided Select-a-Size DNA Clean & Concentrator column (Zymo) and a DNA Clean & Concentrator 5 column (Zymo). A quarter of this reaction was then used for the second round of PCR, in which two different reactions were used to prepare barcode quantification and barcode to variant sequence mapping libraries. For barcode quantification, the second-round amplification was performed using the same primers as used for quantifying cleaved RNA abundance (oJT441 or oJT890 with our Truseq indexing primer) and size selected on an 8% TBE gel. These libraries were sequenced with 50 or 75 bp single end reads on an Illumina NextSeq500. Libraries mapping barcodes to variant sequence were amplified using primers oJT446/447, size selected on an 8% TBE gel, and sequenced with either 75 bp or 150 bp paired end reads on an Illumina MiSeq.

### Barcode to variant sequence mapping and barcode quantification

Barcode quantification and variant sequence identification was handled using custom scripts written in Python. To determine the variant sequence associated with each barcode, the paired end reads from the gDNA-derived libraries were directly parsed as FASTQ files, with one read capturing the barcode and the other the variable region of the construct. Variants with at least one mutation in the constant region proceeding the barcode were discarded. To avoid rare mutations or sequencing errors confounding our variant sequence mapping, variant sequence assignment required at least three reads per barcode. If, among these reads, the second-most frequent sequence was at least 25% as frequent as the top sequence, the barcode was discarded entirely.

Barcodes were counted by directly parsing the single-end sequencing reads derived from both gDNA and RNA in a FASTQ format, with the barcode appearing at a fixed position within the read. As before, any variant with mutations in the constant sequence proceeding the barcode was discarded. Additionally, any two barcodes of identical sequence and unique molecular identifier were counted as a single read. The number of unique reads mapping to each variant barcode was calculated for both the RNA and gDNA-derived samples, and the ratio of RNA to gDNA-derived reads was calculated to extract the relative accumulation of processed mRNA for each variant. Barcodes can then be grouped by associated mutations, thresholded based on read count, and normalized to the median wild-type barcode’s ratio.

## AUTHOR CONTRIBUTIONS

J.T. and G.-W.L. conceptualized project, designed experiments and analysis, and wrote the manuscript with input from all co-authors. J.T. collected data and performed analysis. H.L. and J.D. contributed to the design and implementation of the MPRA experiments. J.-B.L. collected preliminary Rend-seq data for exonuclease knockouts. C.C., S.D., and F. B. constructed strains, provided feedback on analysis, and helped in conceptualization of project.

## ACKNOLWEDGEMENTS

The authors thank David Bechhofer for sharing the phage-cured RNase III knockout strains, Byoung-Mo Koo and Carol Gross for sharing the RNase J2 knockout strain, the MIT BioMicro Center for advice and high-throughput DNA sequencing, Sean McGeary for feedback on early versions of the manuscript, and Michael Laub and David Bartel for feedback in developing this project. Thanks additionally to all members of the Li Lab for valuable discussion. This research is supported by a National Science Foundation Graduate Research Fellowship (to J.C.T.) and the NSF CAREER Award MCB1844668. The C.C. lab is supported by funds from the CNRS (UMR 8261), Université Paris Cité, and the Agence Nationale de la Recherche (CoNoCo and LabEx Dynamo).

---

**Figure S1.**
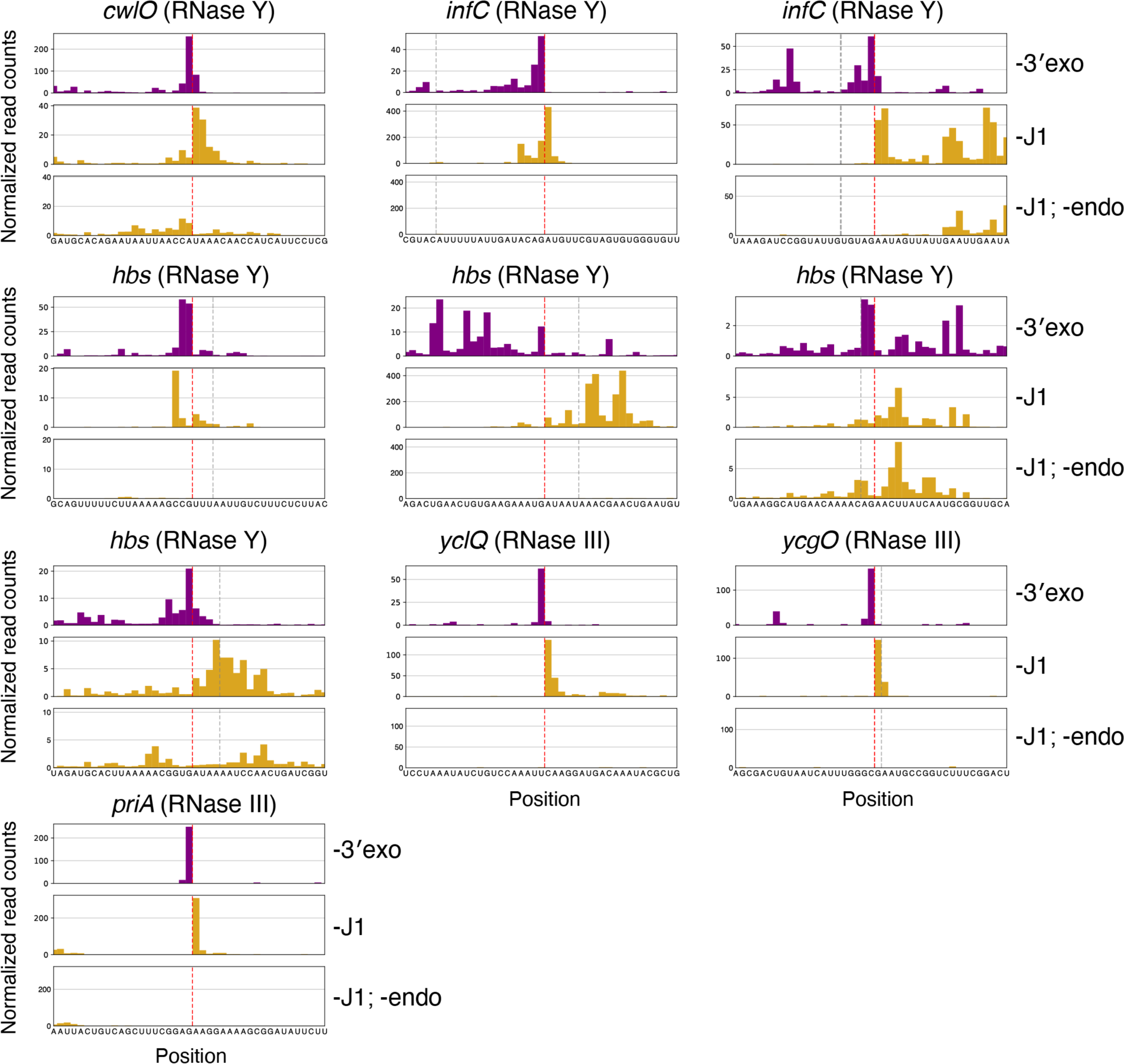
Validation of approach through detection of known positions of endonuclease activity. 5′ and 3′ end sequencing data at known positions of cleavage by RNases Y and III. Datasets considered are a 4-exo knockout (CCB396), *rnjA* knockout (CCB434), and either deletion or depletion of RNase J1 with a knockout of RNase Y (CCB760) or III (BG879). 5′-mapped data shown in yellow and 3′-mapped data shown in purple. Plotted are reads per million CDS-mapping reads, normalized to the average 3′-mapped Rend-seq RPM in this window. A manually annotated position of cleavage based on sequencing data is shown with a red dotted line, and the published position is marked in gray of cleavage if this position disagrees with our annotation. In transcripts with many nearby peak pairs such as *hbs*, additional cleavage positions may be visible beyond that which is highlighted.

**Figure S2.**
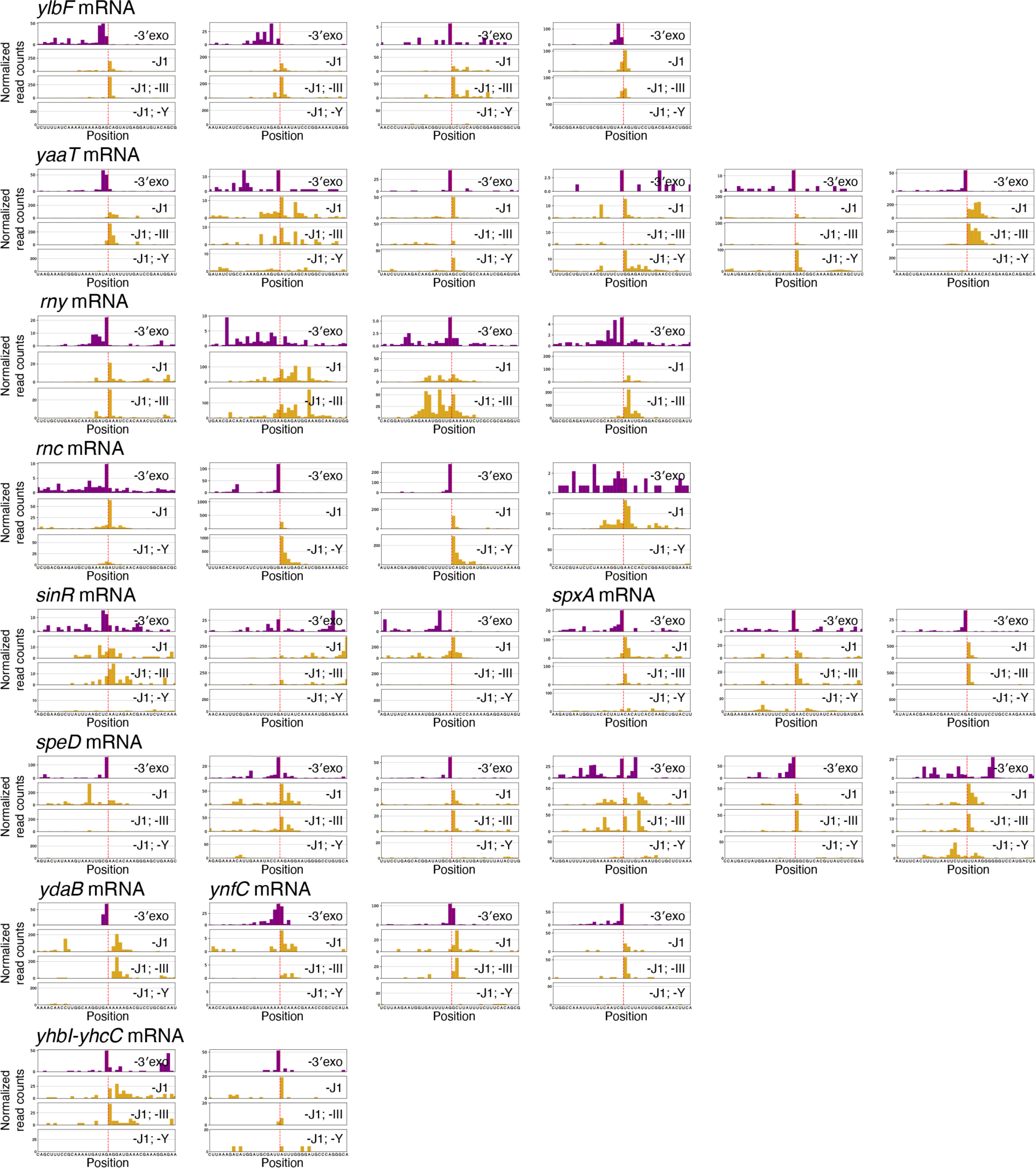
Sequence context and RNase Y/III dependence of cleavage positions within mRNAs known to be destabilized by endoribonucleases. Cleavage positions match order of those shown in Figure 2. Datasets considered are a 4-exo knockout (CCB396), *rnjA* knockout (CCB434), rnjA rnc double knockout in an SPβ and Skin-cured background (BG879), and depletion of RNase J1 with a knockout of *rny* (CCB760). 5′ end sequencing data shown in yellow and 3′ end sequencing data shown in purple. Plotted are reads per million CDS-mapping reads, normalized to the average 3′-mapped Rend-seq RPM in this window. A manually annotated position of cleavage based on sequencing data is shown with a red dotted line. Data from the *rnc*- or *rny*-deficient strain are not shown for sites within the transcripts encoding these genes.

**Figure S3.**
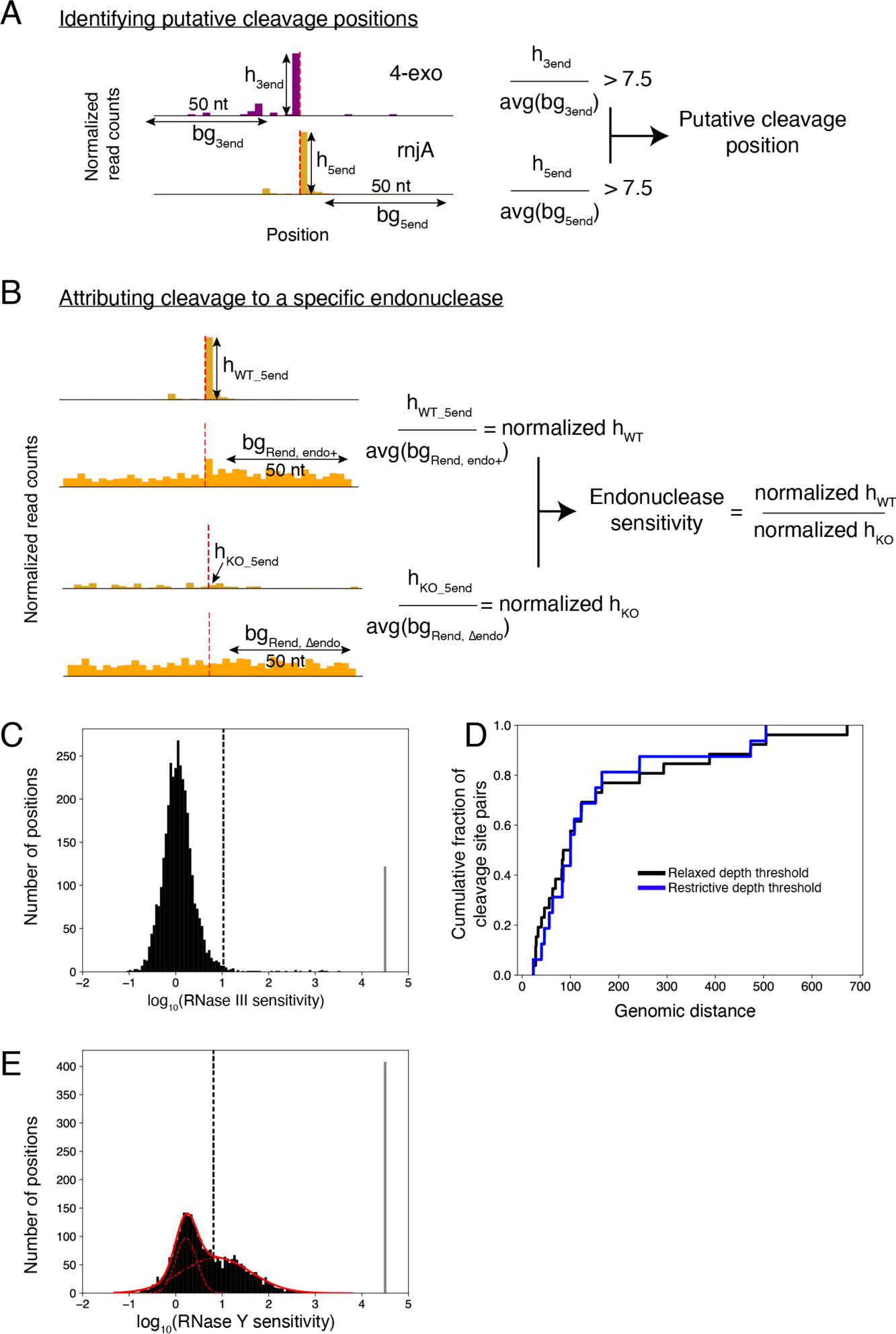
Schematic representation of strategy for calling putative cleavage positions and assigning them to the activity of endoribonucleases. (A) Strategy for calling paired 3′ and 5′ peaks in end sequencing data, with sites exceeding a peak-to-background ratio of 7.5 in the 3′/5′ at adjacent positions called as putative cleavage positions. (B) Strategy for attributing sites to specific endonucleases. Peak heights in 5′ end sequencing are normalized to local density in a corresponding 5′-mapped Rend-seq dataset. A ratio of normalized peak heights between exoribonuclease strains with and without an endoribonuclease knockout (termed “endonuclease sensitivity”) is calculated as a metric for dependence on this endoribonuclease. Note that prior to illustrated calculation, 5′ end sequencing data (h_WT_5end_ and _hKO_5end_) are normalized by the sum of 5′ end sequencing reads mapped to putative cleavage positions, and the Rend-seq background (bg_Rend_) is normalized to the total CDS-mapped reads to account for differences in sequencing depth between samples. (C) Results of systematic identification of RNase III sites with a relaxed 5′ end sequencing depth threshold (as used in Figure 3G, H). Histogram shows distribution of endonuclease sensitivities for called peak pairs in 3′/5′ end sequencing of exoribonuclease knockouts (see Figure S3A and Methods), with black dashed line indicating threshold for calling dependence on RNase III. 162 sites exceeded this defined threshold. For 122 sites we were unable to calculate a sensitivity score due to an absence of 5′ end sequencing counts in our knockout. These sites are called as RNase III sensitive and are counted within the “>4” bin of the histogram. (D) Genomic distance between pairs of unambiguous RNase III cleavage positions identified within 1 kb of one another and predicted to fall on opposite sides of an RNA stem, using both restrictive and relaxed 5′ end sequencing depth thresholds. (E) Results of systematic identification of RNase Y sites with a relaxed 5′ end sequencing depth threshold. Histogram shows distribution of RNase Y sensitivities, plotted as described in Figure 5A. 1351 sites exceeded this defined threshold. For 408 sites we were unable to calculate a sensitivity score due to an absence of 5′ end sequencing counts in our knockout. These sites are called as RNase Y sensitive and are counted within the “>4” bin of the histogram.

**Figure S4.**
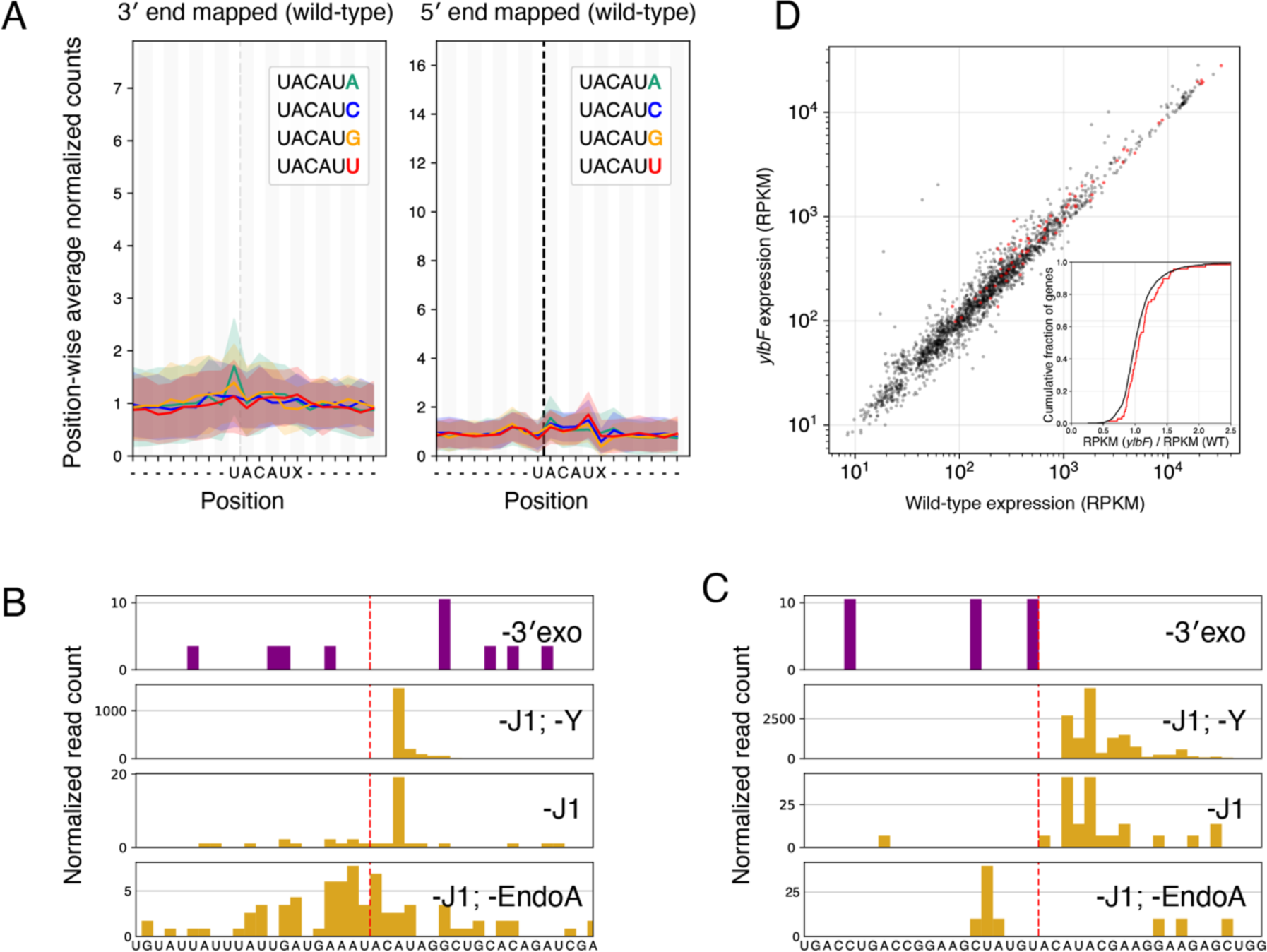
EndoA and YlbF have a limited influence on wild-type *B. subtilis* mRNA decay. (A) UACAUA-specific EndoA cleavage is not detectable in wild-type *B. subtilis.* 3′ (left) and 5′ (right) mapped Rend-seq signal across all UACAU motifs in the genome, separated by downstream nucleotide. Data are derived from wild-type W168. Rend-seq data at each site is normalized to first 8 (for 3′-mapped) or last 8 (for 5′-mapped) positions within the window and a position-wise mean and standard deviation are calculated with 90% winsorization. Motif instances with fewer than 1 read per position or fewer than 10 reads within the normalization window are not considered. Following this filtering, the number of considered sites ranges from 193 to 313 (for 3′-mapped) and 203 to 311 (for 5′-mapped) per motif. The dashed vertical line indicates the position of cleavage by EndoA. (B, C) Representative EndoA sites showing presence (B) or absence (C) of subsequent trimming after the first two nucleotides. Yellow indicates 5′ end sequencing data and purple indicates 3′ end sequencing data. Plotted are reads per million CDS-mapping reads, normalized to the average 3′-mapped Rend-seq RPM in this window. Red dotted line represents manually annotated EndoA cleavage positions. Note that cleavage product accumulation is dependent on EndoA (bottom two panels). (D) mRNA abundances for all annotated *B. subtilis* genes in a wild-type and *ylbF* W168 background. Lowly expressed genes (<128 total reads) are excluded. RPKM calculated as reads per thousand bases of the gene per million total reads that map to *B. subtilis* coding regions. Inset shows differential expression distribution (Δ*ylbF*/WT RPKM) for genes with genes with (red) or without (black) a YlbF-dependent putative cleavage site.

**Figure S5.**
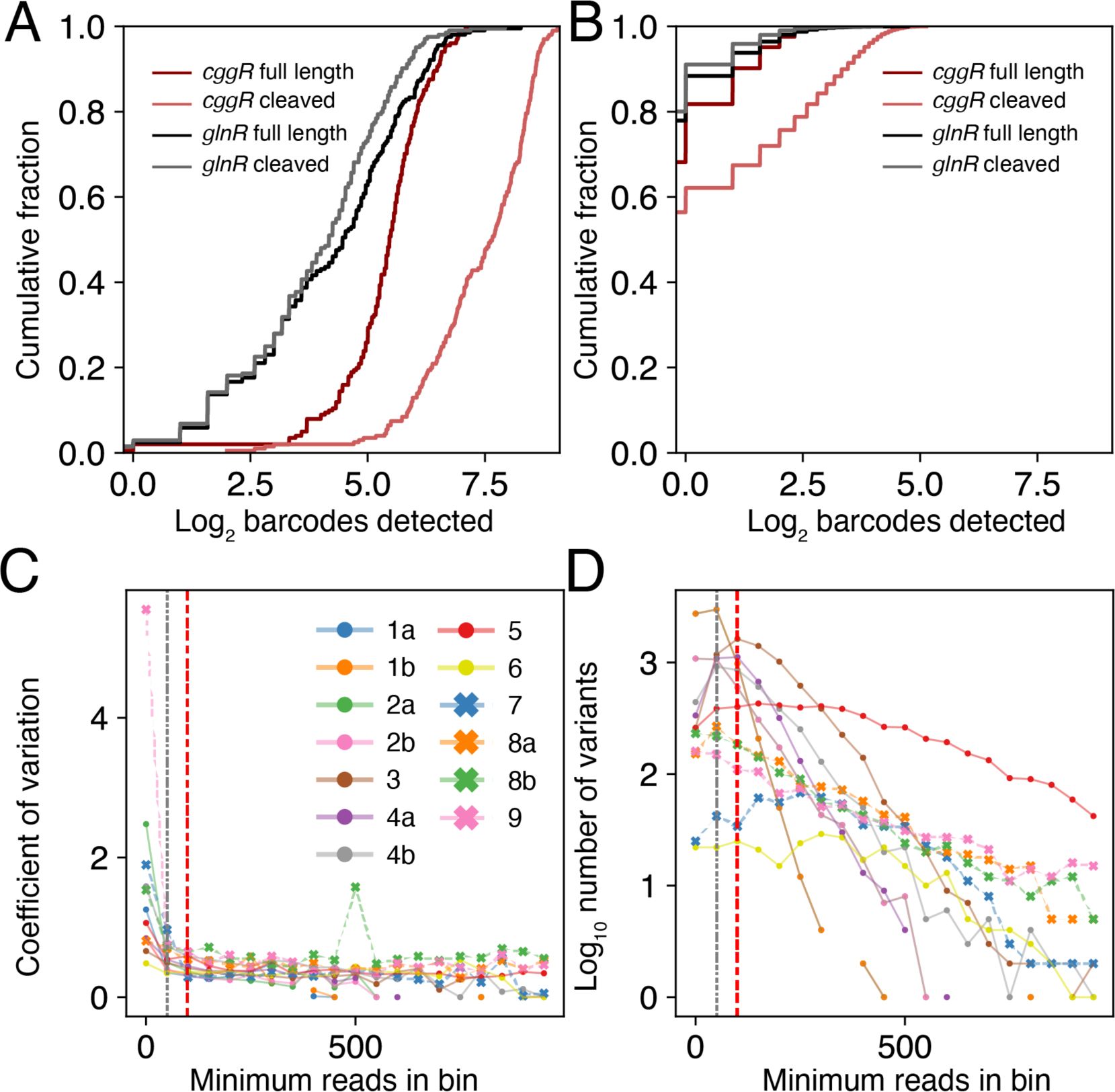
Variability and sequence coverage within our MPRA datasets. (A) Cumulative distribution of the number of unique barcodes associated with each possible sequence containing a single substitution relative to the wild-type *cggR* or *glnR* operon sequence. Undetected sequences are included with a pseudocount of 0.1 prior to log_2_ transformation. (B) Cumulative distribution of the number of unique barcodes associated with each possible sequence containing two substitutions relative to the wild-type *cggR* or *glnR* operon sequence. Undetected sequences are included with a pseudocount of 0.1 prior to log_2_ transformation. (C) Variation in MRPA readout as a function of genomic DNA sequencing depth. The normalized MPRA RNA abundance for each variant with a wild-type sequence was calculated as illustrated in Figure 6A. These values were grouped by the number of unique reads mapping to that variant in the genomic DNA barcode sequencing with a bin width of 50 reads. The coefficient of variation within each group is shown. The gray vertical dashed line indicates the 50 read cutoff used for experiments 4b, 8a, 8b, and 9, red vertical dashed line indicates the 100 read cutoff used in analysis of all remaining experiments. See Table S7 for more information. (D) The number of variants within each bin for each dataset show in (A). Dashed lines show thresholds as described in (A).

**Figure S6.**
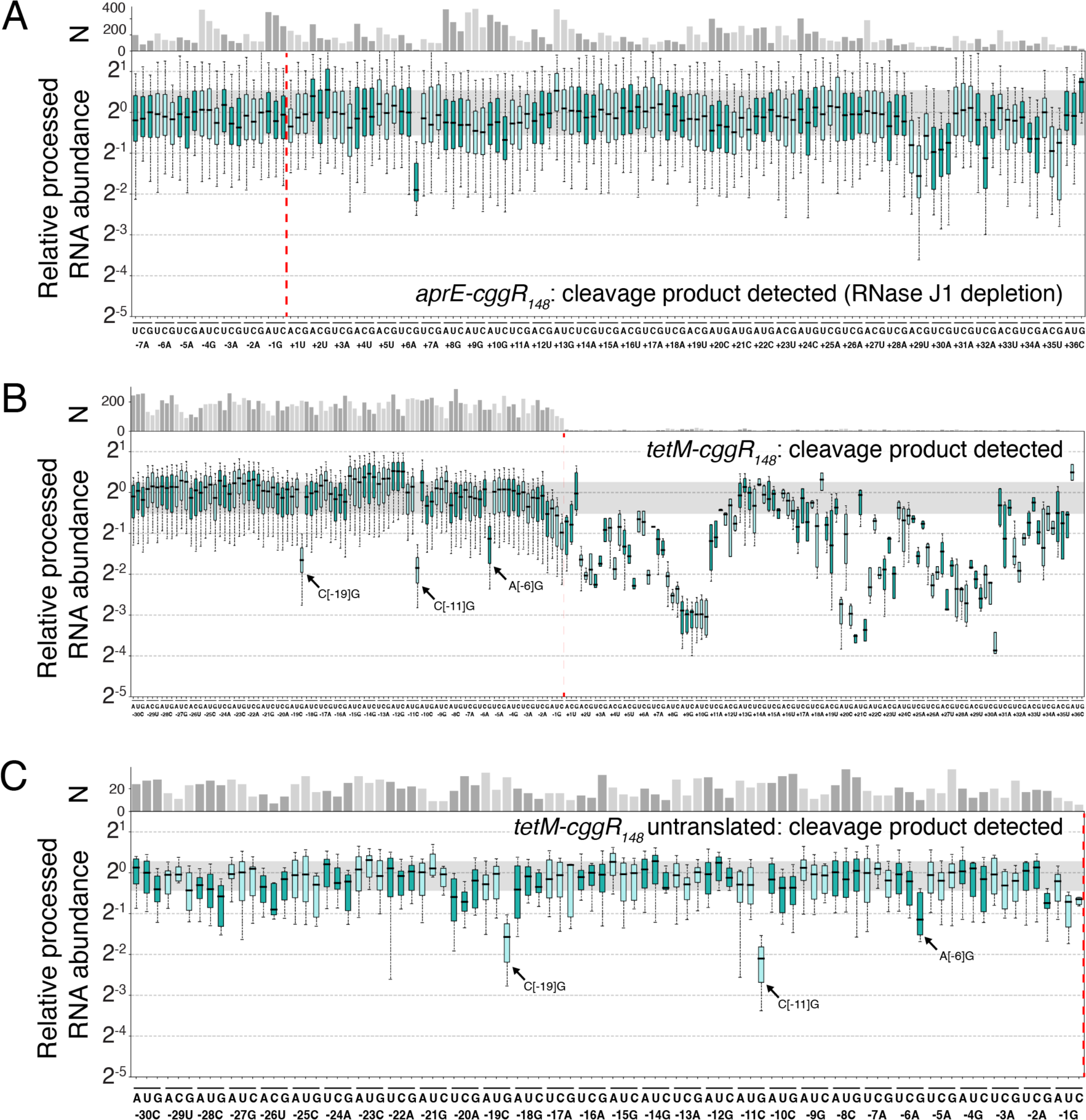
Features driving RNase Y cleavage within *cggR* are consistent across multiple MPRA contexts. (A) Impact of all single-nucleotide mutations on the accumulation of barcoded cleaved *aprE*-*cggR*_148_ RNA in the context of RNase J1 depletion. Boxplots show variation between barcodes of identical variant sequence. Number of barcodes captured for each mutation is indicated above plot. Whiskers indicate 5^th^ and 95^th^ percentile. Gray shaded region indicates interquartile range for variants of wild-type sequence. Red line indicates position of cleavage. Ten variants have a value of zero (no more than one per mutation) and are thus not visualized. (B) Impact of all single-nucleotide mutations on the accumulation of barcoded cleavage product when the *cggR_148_* construct is appended to a new scaffold RNA, *tetM*. Plotted as in (A). Two variants (no more than one per mutation) have a value of zero are thus not visualized. (C) Impact of all single-nucleotide mutations in the 30 nucleotides upstream of the cggR_148_ cleavage site on the accumulation of barcoded cleavage product when a stop codon is inserted upstream of the inserted sequence. *tetM* scaffold was used in this experiment, as in (C). Plotted as in (A).

**Figure S7.**
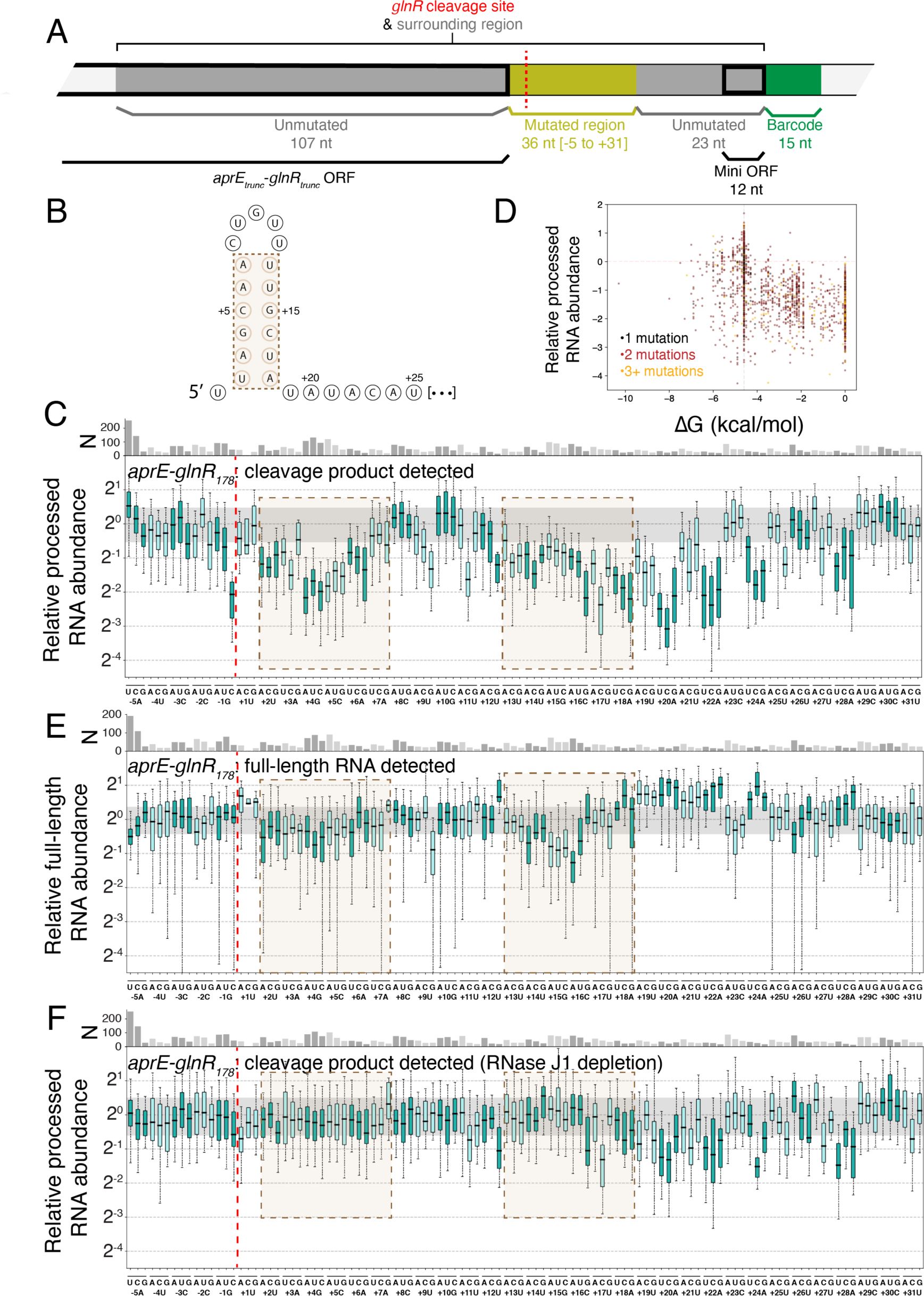
Determinants of *glnR-glnA* mRNA processing by RNase Y. (A) Schematic of the *glnR*_178_ construct inserted into the *aprE* MPRA transcript. *glnR*-derived sequence is colored dark gray, with the mutated positions highlighted in yellow. Translated regions are indicated with a thick border, and the variant barcode is indicated in green. (B) Predicted stem-loop structure at 5′ end of processed RNA. Brown bar corresponds to regions highlighted in panels C, E, and F. (C) Impact of all single-nucleotide mutations on the accumulation of barcoded cleaved RNA. Boxplots show variation between barcodes of identical variant sequence. Number of barcodes captured for each mutation is indicated above plot. Whiskers indicate 5^th^ and 95^th^ percentile. Gray shaded region indicates interquartile range for variants of wild-type sequence. Red line indicates position of cleavage. Brown bar corresponds to positions predicted to form the stem of the structure showed in (B). Nine variants (no more than two per mutation) have a value of zero are thus not visualized. (D) Relationship between predicted strength of downstream secondary structure and accumulation of barcoded cleavage product. All data are derived from experiment 8a (Table S7). The vertical dashed line indicates the ΔG of the unmutated sequence. 17 variants fall outside of the bounds of this plot. (E) Impact of all single-nucleotide mutations on the accumulation of barcoded full-length RNA, plotted as described in (C). 30 variants (no more than two per mutation) have a value of zero are thus not visualized. (F) Impact of all single-nucleotide mutations on the accumulation of barcoded cleaved RNA in the context of RNase J1 depletion, plotted as described in (C). Three variants (no more than one per mutation) have a value of zero are thus not visualized.

**Table S1. List of strains used in this study**

**Table S2. List of oligonucleotide sequences used in this study**

**Table S3. List of identified RNase III cleavage sites**

**Table S4. List of identified RNase Y cleavage sites**

**Table S5. List of putative RNase III cleavage sites with relaxed 5’ end sequencing depth threshold**

**Table S6. List of putative RNase Y cleavage sites with relaxed 5’ end sequencing depth threshold**

**Table S7. Summary of MPRA experiments conducted in this study**

